# Multiomics reveals age-dependent metabolic reprogramming of macrophages by wound bed niche secreted signals

**DOI:** 10.1101/2024.10.30.621159

**Authors:** Maria Fernanda Forni, Gabriela A. Pizzurro, Will Krause, Amanda F. Alexander, Kate Bridges, Yiting Xu, Olivia Justynski, Anthony Gabry, Niels Olsen Saraiva Camara, Kathryn Miller-Jensen, Valerie Horsley

## Abstract

The cellular metabolism of macrophages depends on tissue niches and can control macrophage inflammatory or resolving phenotypes. Yet, the identity of signals within tissue niches that control macrophage metabolism is not well understood. Here, using single-cell RNA sequencing of macrophages in early mouse wounds, we find that, rather than gene expression of canonical inflammatory or resolving polarization markers, metabolic gene expression defines distinct populations of early wound macrophages. Single-cell secretomics and transcriptomics identify inflammatory and resolving cytokines expressed by early wound macrophages, and we show that these signals drive metabolic inputs and mitochondrial metabolism in an age-dependent manner. We show that aging alters the metabolome of early wound macrophages and rewires their metabolism from mitochondria to glycolysis. We further show that macrophage-derived Chi3l3 and IGF-1 can induce metabolic inputs and mitochondrial mass/metabolism in aged and bone marrow-derived macrophages. Together, these findings reveal that macrophage-derived signals drive the mitochondrial metabolism of macrophages within early wounds in an age-dependent manner and have implications for inflammatory diseases, chronic injuries, and age-related inflammatory diseases.

**In Brief:** This study reveals that macrophage subsets in early inflammatory stages of skin wound healing are defined by their metabolic profiles rather than polarization phenotype. Using single-cell secretomics, we establish key macrophage cytokines that comprise the *in vivo* wound niche and drive mitochondrial-based metabolism. Aging significantly alters macrophage heterogeneity and increases glycolytic metabolism, which can be restored to OxPHOS-based metabolism with young niche cytokines. These findings highlight the importance of the tissue niche in driving macrophage phenotypes, with implications for aging-related impairments in wound healing.

**Highlights:**

- Single cell transcriptional analysis reveals that reveals that metabolic gene expression identifies distinct macrophage populations in early skin wounds.
- Single-cell secretomic data show that young macrophages contribute to the wound bed niche by secreting molecules such as IGF-1 and Chi3l3.
- Old wound macrophages display altered metabolomics, elevated glycolytic metabolism and glucose uptake, and reduced lipid uptake and mitochondrial mass/metabolism.
- Chi3l3 but not IGF-1 secretion is altered in macrophages in an age dependent manner.
- Chi3l3 can restore mitochondrial mass/metabolism in aged macrophages.

## Introduction

Restoration of tissue function after damage induced by disease or injury is essential for life. In most mammalian tissues, tissue repair occurs in two phases: an early inflammatory response that clears microbes, pathogens, and cellular debris and a resolution process that regenerates specific cell types with or without a scar, depending on the damage and tissue type. Both of these phases of wound healing can be regulated by macrophages, which can adopt a range of inflammatory or resolving phenotypes (Krzyszczyk et al., 2018). When dysfunctional, macrophage activity or phenotypes are associated with inflammatory diseases, cancer, or chronic injuries (Azizi et al., 2018). Much of our understanding of macrophage phenotypes comes from *in vitro* models in which macrophages derived from bone marrow are stimulated with inflammatory cues like IFNg and/or LPS or a resolving cytokine, IL-4, to produce polarized bone marrow-derived macrophages (BMDMs). Yet, within tissues, macrophage phenotypes depend on the specific niche, and tissue macrophage heterogeneity cannot be fully recapitulated in ex vivo (Pang et al., 2022). Thus, defining the mechanisms that control the transition of macrophages from an inflammatory to a resolving phenotype within the complex and dynamic environment of damaged tissues is paramount.

The skin is an excellent model for defining the mechanisms that control macrophage transitions after injury due to the well-defined phenotypic and temporal evolution of macrophages from inflammation to resolution after a mechanical breach of the skin’s barrier. Initially, damage recruits monocytes to the skin in a CCR2-dependent manner, generating type-1-activated pro-inflammatory and pro-angiogenic Ly6Chi+CD11b+F480+ macrophages (Pang et al., 2022). As healing ensues, macrophages transition toward a type-2-activated resolution phenotype associated with changes in the expression of cytokine mRNAs and expression of mannose receptor (CD206), CD301b, Fizz1, IL-10, transforming growth factor (TGF)-β1, and vascular endothelial growth factor (VEGF) (Ferrante & Leibovich, 2012; Mirza et al., 2009; Shook et al., 2016) to support repair of keratinocytes to restore the skin’s barrier and fibroblasts, endothelial and other cell types.

Metabolic regulation of immune cells is an emerging theme that is starting to be understood within tissues (Eming et al., 2021). *In vitro* studies have shown that IFNg/LPS polarized inflammatory BMDMs rely on glycolysis and the cyclic pentose phosphate pathway, not mitochondrial metabolism, to generate inflammatory cytokines (Bala et al., 2021). Conversely, IL-4-induced resolving BMDMs catabolize lipid molecules via the TCA cycle and use oxidative phosphorylation to generate ATP and suppress inflammatory genes (Oishi et al., 2017). A recent study revealed that oxidative phosphorylation in early-stage, inflammatory skin macrophages promote angiogenesis and inflammation and identified subpopulations of macrophages that displayed distinct mRNA expression of metabolic genes (Willenborg et al., 2021). Thus, metabolic regulation of polarized macrophages *in vivo* is driven by the injured tissue niche in a distinct manner from metabolic regulation of BMDM phenotypes.

Here, by comparing polarized BMDMs and early mouse wound macrophages via RNA sequencing and secretomics at the single-cell level, we find that, unlike inflammatory or resolving BMDMs, macrophages in mouse inflammatory skin wounds display a heterogeneous mix of inflammatory and resolving mRNA expression and cytokine production. Interestingly, subpopulations of wound-derived macrophages can be defined by metabolic gene expression and display distinct metabolic profiles compared to polarized BMDMs, and metabolite production and metabolic pathway use are altered in early wound macrophages with age. Characterization of cytokines within inflammatory mouse wounds and secreted by wound-derived macrophages revealed that Chi3l3 and IGF-1 are key tissue niche factors that can change the preferential energetic substrates and metabolism of macrophages and rescue metabolic defects associated with age. Taken together, our data indicate that macrophage-derived cytokines rewire macrophage metabolism in an age-dependent manner.

## Results

### Single-cell analysis reveals that macrophage populations are heterogeneous, displaying pro and anti-inflammatory characteristics in early wound beds

To better understand how the heterogeneity of the early wound bed macrophages (**Figs. 1A and S1**) compares to the classic polarization schemes that induce pro-inflammatory and resolving macrophage phenotypes *in vitro,* we compared single cell RNA sequencing (scRNA-seq) data from early wounds of mouse skin (**Fig. 1A-B**) or from BMDMs left untreated or treated with LPS+IFN-γ (pro-inflammatory) or IL-4 (resolving) for 6 hours (**Fig. 1C-D**) (Muñoz-Rojas et al., 2021, Justynski et al., 2023). Visualization of BMDM scRNA-seq data with Uniform Manifold Approximation and Projection (UMAP) (Becht et al., 2018, 2019) revealed complete separation between the three treatments separate in the two-dimensional space, suggesting that LPS+IFN-γ and IL-4 induce distinct transcriptional states (**Fig. 1D**). As expected, macrophages treated with LPS+IFN-γ expressed canonical pro-inflammatory-associated genes (including *Il1b, Nos2, Tnf, Il6*, and *Cd86*), while macrophages treated with IL-4 expressed traditional resolving or anti-inflammatory-associated genes (including *Arg1*, *Retnla,* and *Chil3*) (**Figs. 1D-E**).

**Figure 1.**
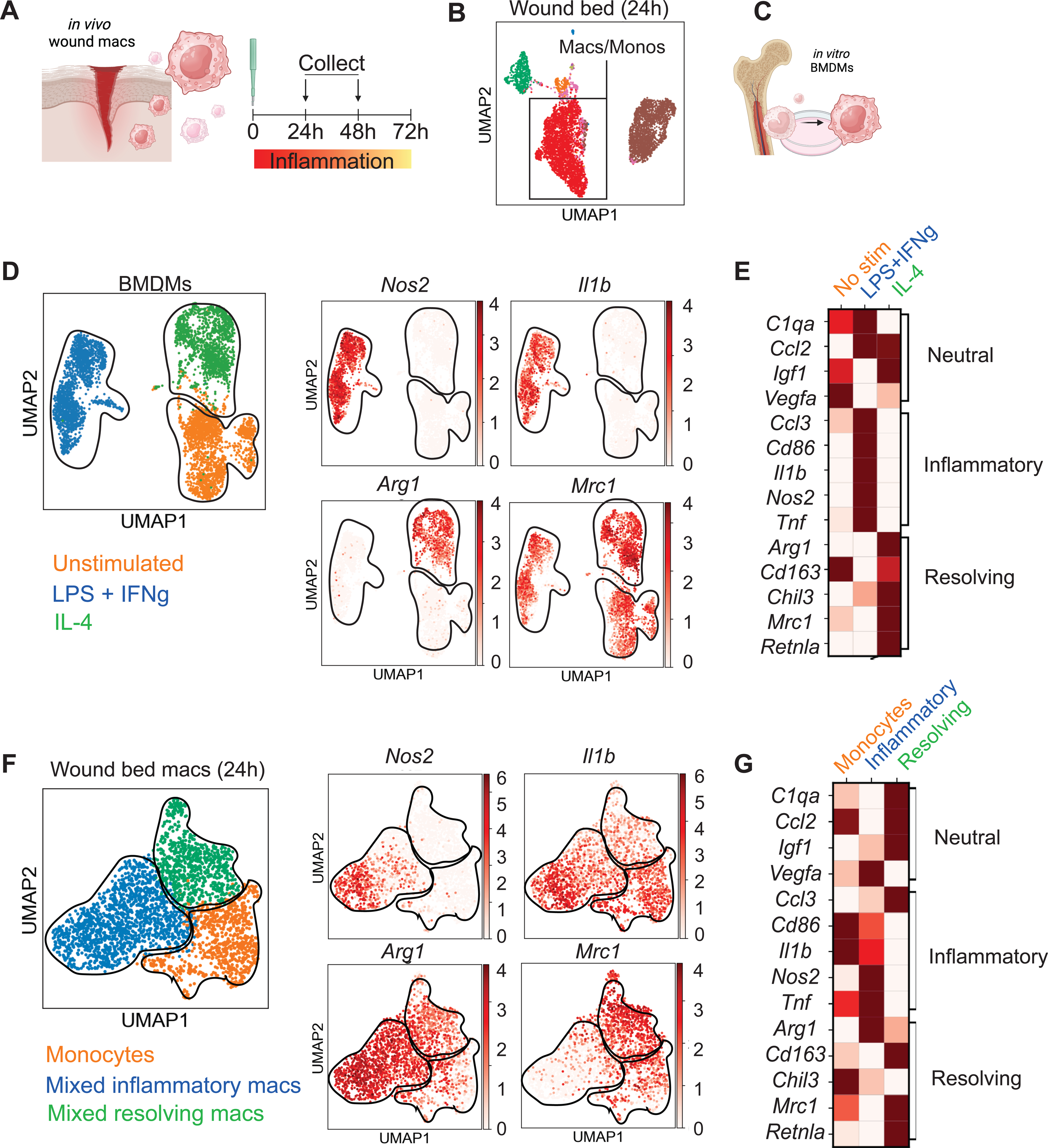
Single-cell RNA sequencing reveals distinct populations of macrophages in early skin wounds compared to polarized bone marrow-derived macrophages. A. Schematic of isolation of wound macrophages in 24 hour (h) and 48h wound beds. B. UMAP plots of scRNA sequencing data of wound bed cells. Data are from GSE223660. C. Schematic and UMAP plot of scRNA sequencing data from unstimulated BMDMs or BMDMs stimulated with IFN/LPS or IL-4. D. UMAP and feature plots of scRNA sequencing data from unstimulated BMDMs or BMDMs stimulated with IFN/LPS or IL-4. E. Heatmap of mRNAs associated with Neutral, Inflammatory, and Resolving macrophages in BMDM scRNA sequencing data. F. UMAP and feature plots of scRNA sequencing data from 24-hour (h) macrophages (macs). G. Heatmap of mRNAs associated with neutral, inflammatory, and resolving macrophages in 24h wound bed macrophage scRNA sequencing data.

Next, we examined macrophage heterogeneity in mouse skin wounds *in vivo*. To this end, we analyzed macrophages within early skin wounds 1 and 2 days after injury, when cells are generally considered pro-inflammatory (Justynski et al., 2023). Unbiased Leiden clustering of myeloid cells *Itgam*+ and *Adgre1*+ cells revealed three distinct populations of macrophages in early 24 hour wounds (**Fig. 1F**). We first plotted the scRNA-seq data with a diffusion mapping dimensionality reduction algorithm PHATE (potential of heat diffusion for affinity-based transition embedding) (Moon et al., 2019; Moon et al., 2020) to emphasize relational trajectories between cells (**Fig. S1A**). We then used RNA velocity to infer how cell transcription changes with time (**Fig. S1B**) (La Manno et al., 2018). We found that cluster 0, which co-expressed *Ly6c2* and *Ccr2* characteristic of monocytes, split into two macrophage branches in the wound bed (expressing both *Adgre1* and *Csf1r*) (**Fig. S1C**). We also noted a small population of macrophages expressing *Lyve1*, which could indicate tissue resident macrophages at this early time point (Justynski et al., 2023).

To determine the phenotype of the macrophage clusters, we replotted the clusters using UMAP and examined the gene expression of canonical pro-inflammatory and resolving macrophage markers (**Fig. 1F**). One cluster expressed high levels of the classic inflammation-associated genes, including *Il1b, Nos2, Tnf,* and *Cd86*. However, this cluster of Inflammatory macrophages also expressed high levels of resolving genes, including *Vegfa* and *Arg1*, and thus we refer to this cluster as ‘mixed Inflammatory’. The other macrophage cluster displayed high levels of several genes associated with an anti-inflammatory or resolving state (*Mrc1*, *Retnla,* and *Cd163*). However, macrophages in this cluster also expressed inflammatory genes including *Ccl3* and significantly less *Arg1* and *Chil3* than other clusters, and so we refer to them as ‘mixed resolving’ (**Fig. 1G**). Interestingly, the monocyte cluster expressed high levels of *Ccl2*, but also *Chil3* and *Il-1b*. Notably, by 48h post-wounding, the monocytes appeared to have largely differentiated into the two macrophage clusters (**Fig. S1D-E**). Thus, unlike BMDM macrophages differentiated from bone marrow *in vitro*, early mouse wounds *in vivo* display diverse, heterogeneous populations of macrophages with mixed inflammatory and resolving transcriptomes..

### The heterogeneity of macrophage populations in early wound beds is a direct effect of the wound niche and is absent in naive skin

To evaluate macrophage heterogeneity in early wound beds at the protein level, we performed high-dimensional flow cytometry analysis of cells isolated from naive skin or 24h wounds with a panel of 15 markers expressed by inflammatory or resolving skin macrophage populations (**Fig. 2A, S1 and S5**). We visualized the relationships between protein expression of single cells using UMAP plots and found that macrophages in naive skin have little overlap with wound-derived macrophages (**Fig. 2B**).

**Figure 2.**
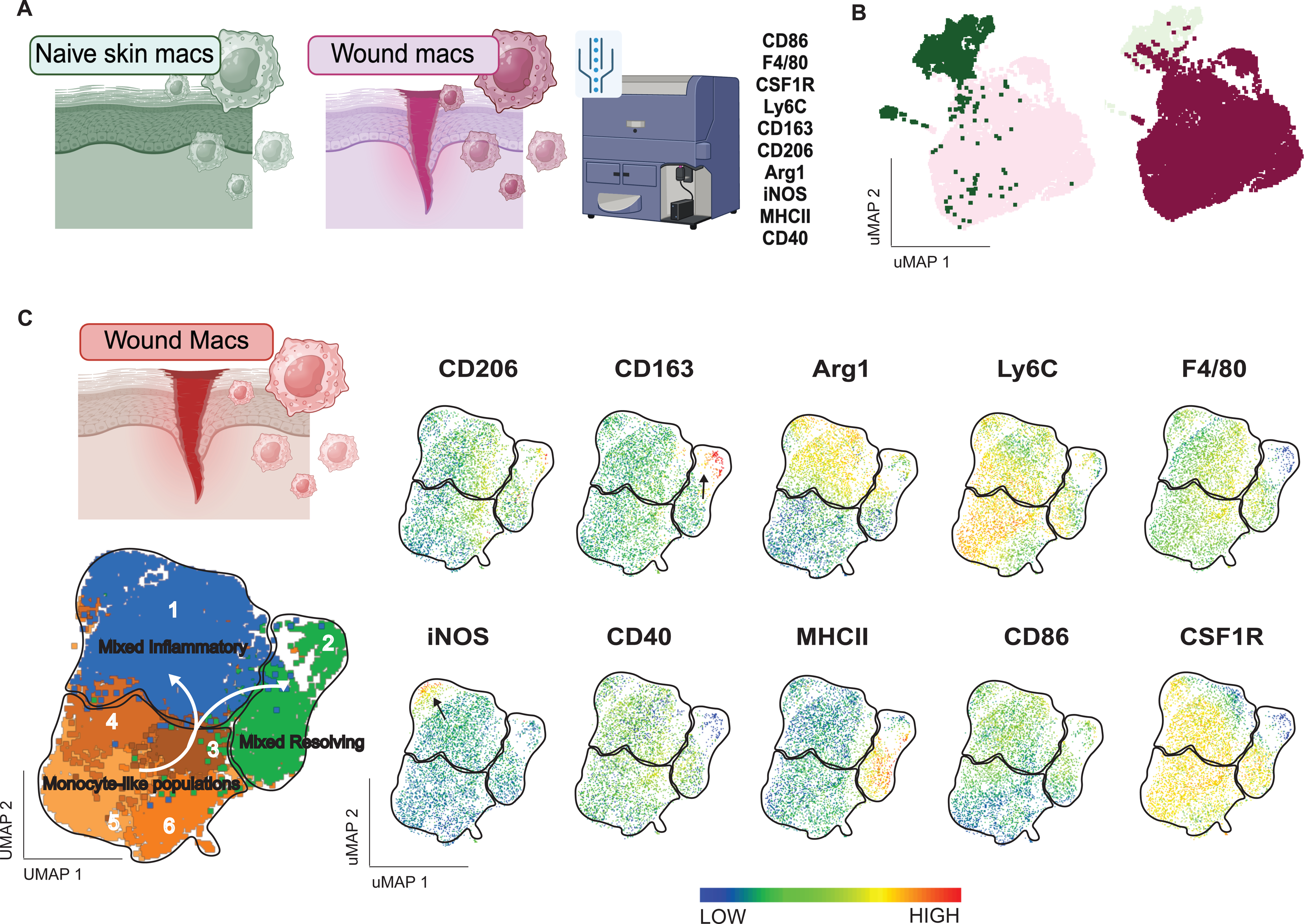
High dimensional flow cytometry analysis of 24h wound bed macrophages. A. Schematic of isolation of naive and 24h wound macrophages and analysis of neutral (green), inflammatory (orange), and resolving (blue) markers. B. UMAP plots of flow cytometry data of naive and wound bed macrophages. Data from x wound beds of y mice were combined. C. Schematic and UMAP and feature plots of flow cytometry data from 24h wound beds.

Next, we visualized the wound macrophages separately and identified three broad cell clusters. All clusters expressed relatively uniform levels of CSF1R, while clusters 1 and 2 expressed slightly more F4/80 (consistent with differentiated macrophages) and cluster 3 (comprised of 4 small clusters) was enriched for LY6C (**Fig. 2C**), consistent with monocytes. Cluster 1 expressed a subpopulation of cells that were high for the inflammatory marker iNOS, but also expressed high levels of resolving proteins such as Arg1 and CD206, suggesting a mixed inflammatory phenotype. Cluster 2 expressed a subpopulation with high levels of the resolving protein CD163, but also expressed high levels of the inflammatory proteins MHCII and CD86, indicating a mixed resolving phenotype (**Fig. 2C**). These data indicate that even at the protein level, macrophages within early stage wound beds express a mix of proteins associated with inflammatory and resolving phenotypes.

### Different metabolic profiles distinguish *in vitro* and *in vivo* macrophage subsets

Inflammatory BMDMs are known to primarily utilize sugars via glycolysis, while resolving BMDMs metabolize lipids via fatty acid oxidation (Eming et al., 2021). We were therefore interested in whether our data supports a link between tissue-specific macrophages phenotypes and their metabolic state. To do this, we examined the macrophage-specific expression of metabolic genes in the scRNA-seq data from BMDMs and early wounds by scoring each single cell by its average expression of genes associated with glycolysis, TCA cycle, and lipid metabolism (**Fig3A** and **Supplementary Table 1**). Surprisingly, metabolic gene signatures were not specific to any BMDM polarization state (**Fig. 3B**). By contrast, specific metabolic signatures were enriched in particular macrophage clusters from 24h mouse wound beds. The mixed inflammatory population was enriched for expression of genes associated with glycolysis and the TCA cycle, while the mixed resolving macrophage cluster was enriched for expression of genes associated with lipid metabolism (**Fig. 3B-C**).

**Figure 3.**
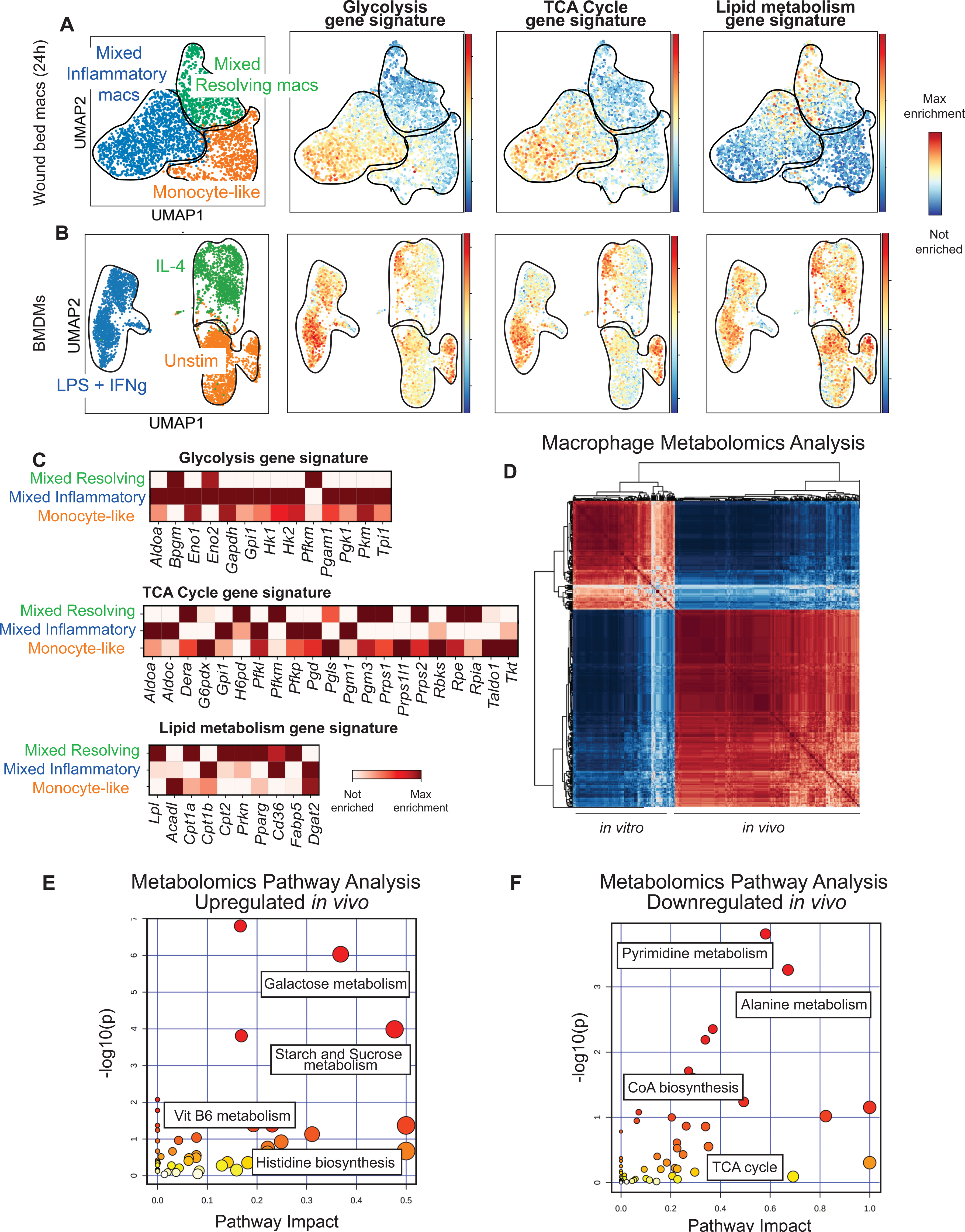
Metabolic gene expression and metabolomics of early wound bed macrophages. A-B. UMAP plots of BMDMs or 24h mouse wound bed macrophages indicating metabolic signatures for glycolysis, TCA cycle, and lipid metabolism genes. C. Heat maps of metabolic genes in BMDMs. D. Heat map of metabolic analysis of IFN/LPS stimulated polarized macrophages and 24h wound bed macrophages. E-F. Pathway analysis plots for metabolism pathways upregulated or downregulated in 24 -hour wound bed macrophages.

Next, we analyzed metabolic output of LPS+IFNg polarized BMDMs and 24h wound-derived F4/80+ macrophages with high-resolution targeted metabolomics to further analyze the metabolic changes associated with macrophages with different phenotypes. A total of 2,322 unique metabolites were annotated using mass spectrometry, and metabolite abundance was determined after standardization and normalization. Comparing metabolite abundance between BMDMs and 24h wound beds macrophages revealed 252 metabolites that were significantly different between BMDMs and wound bed-derived macrophages (**Fig. 3D**). Metabolic pathway analysis showed that macrophages in 24h wound beds upregulated glucose and starch metabolites, galactose-related metabolites, and histidine metabolism precursors (**Fig. 3E**), whereas TCA-related metabolites were downregulated compared to BMDMs (**Fig. 3F**). These data confirm that polarized BMDMs have distinct phenotypes compared to 24h wound bed-derived macrophages, and suggest that metabolic pathways may regulate the different phenotypes of mixed inflammatory vs. resolving macrophages within mouse skin wounds.

### Early wound bed macrophages display a heterogeneous secretion profile containing inflammatory and resolving cytokines

We hypothesized that differences between BMDMs and wound macrophages might arise from signals within the complex wound environment. To examine which ligands are present within 24h wounds of mouse skin, we analyzed the expression of more than 40 ligands or secreted molecules that function in tissue repair (**Table S2**) in scRNA-seq data from 24h wounds (**Fig. 4A**). Fibroblasts and neutrophils expressed the highest number of mRNAs for the selected ligands, and macrophages expressed high levels of both inflammatory and resolving mRNAs, including *Vegfa*, *Chil3*, *Ccl2*, and *Il1b* (**Fig. 4A**).

**Figure 4.**
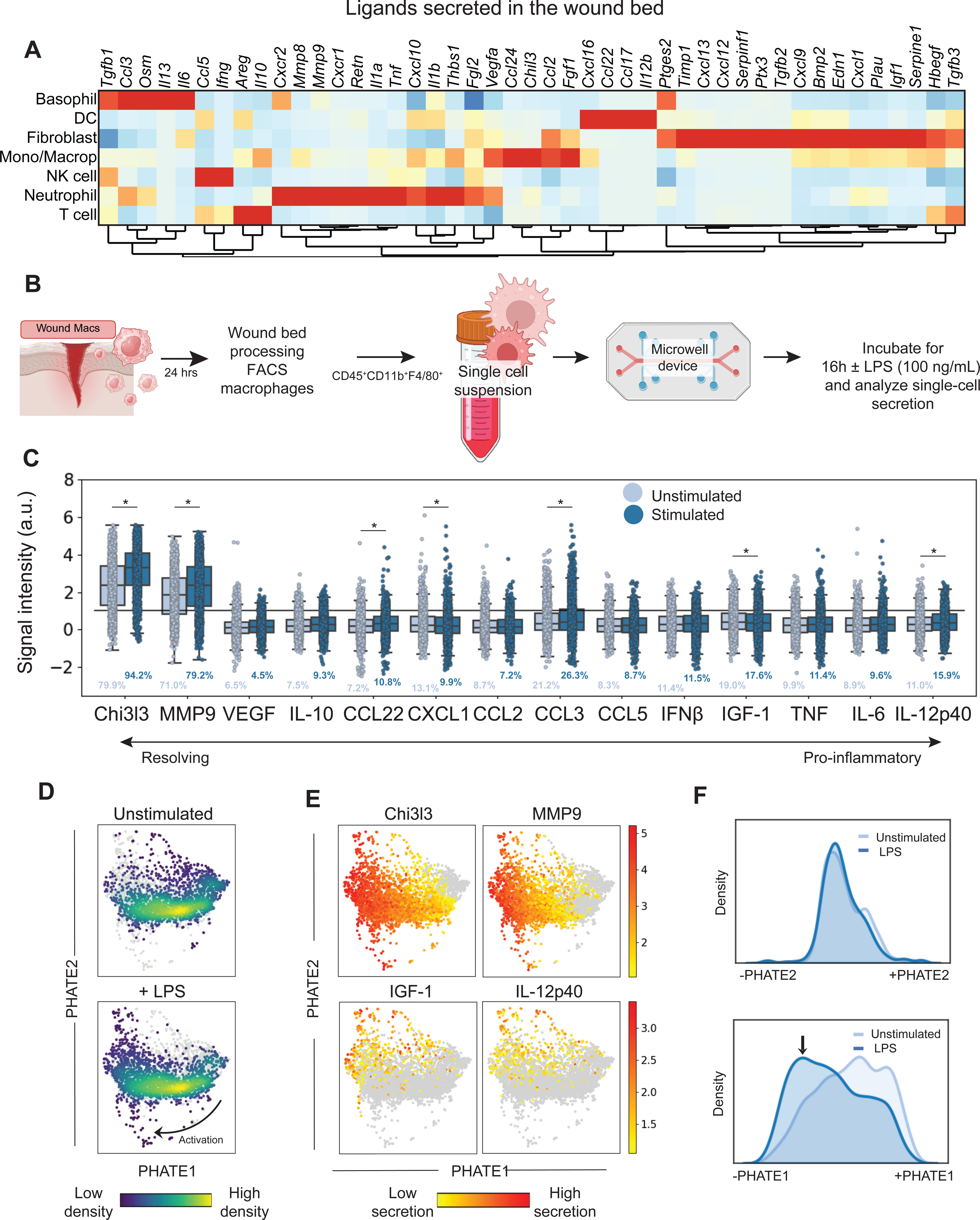
Single-cell RNA sequencing and secretion analysis reveals stable mixed phenotype of 24h wound bed macrophages. A. Heatmap of gene expression in 24 wounds for ligands expressed by indicated cell types. B. Schematic illustrating the experimental design for single-cell secretomic analysis. C. Box and whisker plots of single-cell secretion from F4/80+ macrophages isolated from 24h mouse wounds with or without LPS stimulation. D-E. 2D PHATE visualization of x-dimensional single-cell secretion from F4/80+ macrophages isolated from 24h mouse wounds either unstimulated or treated with LPS. Data include macrophages that secrete at least one measured cytokine above the threshold colored by (D) the kernel density estimation (KDE) for unstimulated or LPS-treated macrophages with grey indicating the other condition or (E) relative intensity of the indicated cytokine (cells lacking secretion in grey). F. Kernel density estimation (KDE) for individual F4/80+ macrophages isolated from 24h mouse wounds along the indicated PHATE axis calculated from data in (D).

Since macrophages were one of the most prevalent cell types in early mouse wounds (**Fig1B**) (Justynski et al., 2023), we further hypothesized that macrophages influence their heterogeneity through autocrine and paracrine signaling. To evaluate the secretory heterogeneity of different macrophages in early wounds, we performed multiplexed single-cell secretion profiling of macrophages using our single-cell secretome microwell assay (**Fig. 4A**) (Lu et al., 2013; Munoz-Rojas et al., 2021; Muñoz-Rojas et al., 2021). We FACS purified live, CD45+, F4/80+, and CD11b+ macrophages from 24h wound beds, cultured them in microwells for 16h, and measured the concentration of 15 different ligands. Surprisingly, most early wound-derived macrophages secreted high levels of proteins associated with resolving macrophages, including Chi3l3 (the protein product of *Chil3*) and matrix metalloproteinase 9 (MMP9), while much smaller fractions secreted inflammatory proteins like TNF and CXCL1 (**Fig. 4B and 4C**).

To determine if polarizing wound bed macrophages to a more inflammatory phenotype could enhance their production of inflammatory cytokines, we profiled protein secretion after 16h of LPS stimulation, which we found previously could alter BMDM secretion patterns (Alexander et al., 2021). LPS-stimulated macrophages displayed significantly altered secretion of a range of resolving and inflammatory cytokines: Chi3l3, MMP9, CCL22, CXCL1, CCL3, and IGF-1 (**Fig. 4C**), instead of alterations in solely inflammatory proteins.

To further explore how LPS could globally alter the single-cell secretion patterns, we visualized our single-cell secretion profiling data in two dimensions using PHATE (**Fig. 4D**). LPS-stimulation induced wound bed macrophages to secrete a mixture of inflammatory and resolving cytokines, including Chi3l3, MMP9, IGF-1, and IL-12p40 expression (**Fig. 4E)**. PHATE analysis revealed that wound-derived macrophages lie along a continuum in two-dimensional space, where activation by LPS stimulation was represented along the negative PHATE1 axis (**Fig. 4F**), consistent with our prior data on the impact of LPS stimulation on BMDMs (Alexander et al., 2021). In particular, secretion of Chi3l3 and MMP9 were strongly correlated with the PHATE1 axis, while other proteins displayed more modest correlations (**Fig. S2)**.

### Aging rewires the secretome of early wound bed macrophages

Since aged macrophages display defective responses to polarizing signaling, wound healing capabilities and reduced plasticity (Ding et al., 2021), we examined how macrophage heterogeneity and phenotypes were altered by performing high-dimensional flow cytometry of macrophages in in 24h mouse wounds of 48-month-old mice (**Fig. 5A**). As in the young mice (**Fig. 2C**), we identified three broad macrophage clusters in wounds of aged mice. While all clusters expressed markers CSF1R and F4/80, several clusters (4, 5, and 6) expressed high levels of Ly6C, indicating they are monocyte-like cells (**Figs. 5A-B**). Cluster 1 in aged mouse wounds expressed moderately high levels of moderately high levels of iNOS and MHCII (which are thought to be inflammatory proteins) (**Figs. 5A-B)**, but did display a subpopulation with high levels of iNOS as that we observed in young mice (**Fig. 2C**). This cluster also expressed more Arg1 than other clusters and expressed CD206 as well as MHCII, suggesting a mixed inflammatory phenotype. Cluster 2 expressed Arg1 and CD206 and high levels of MHCII (**Figs. 5A-B)**, which is similar to the mixed resolving population in young wound macrophages (**Fig. 2A**). However, overall, these multidimensional FACS data reveal that aged wound macrophages have less diversity in their protein expression of these markers (**Figs. 5A-B**) compared to macrophages from young wounds (**Fig. 2A**).

**Figure 5.**
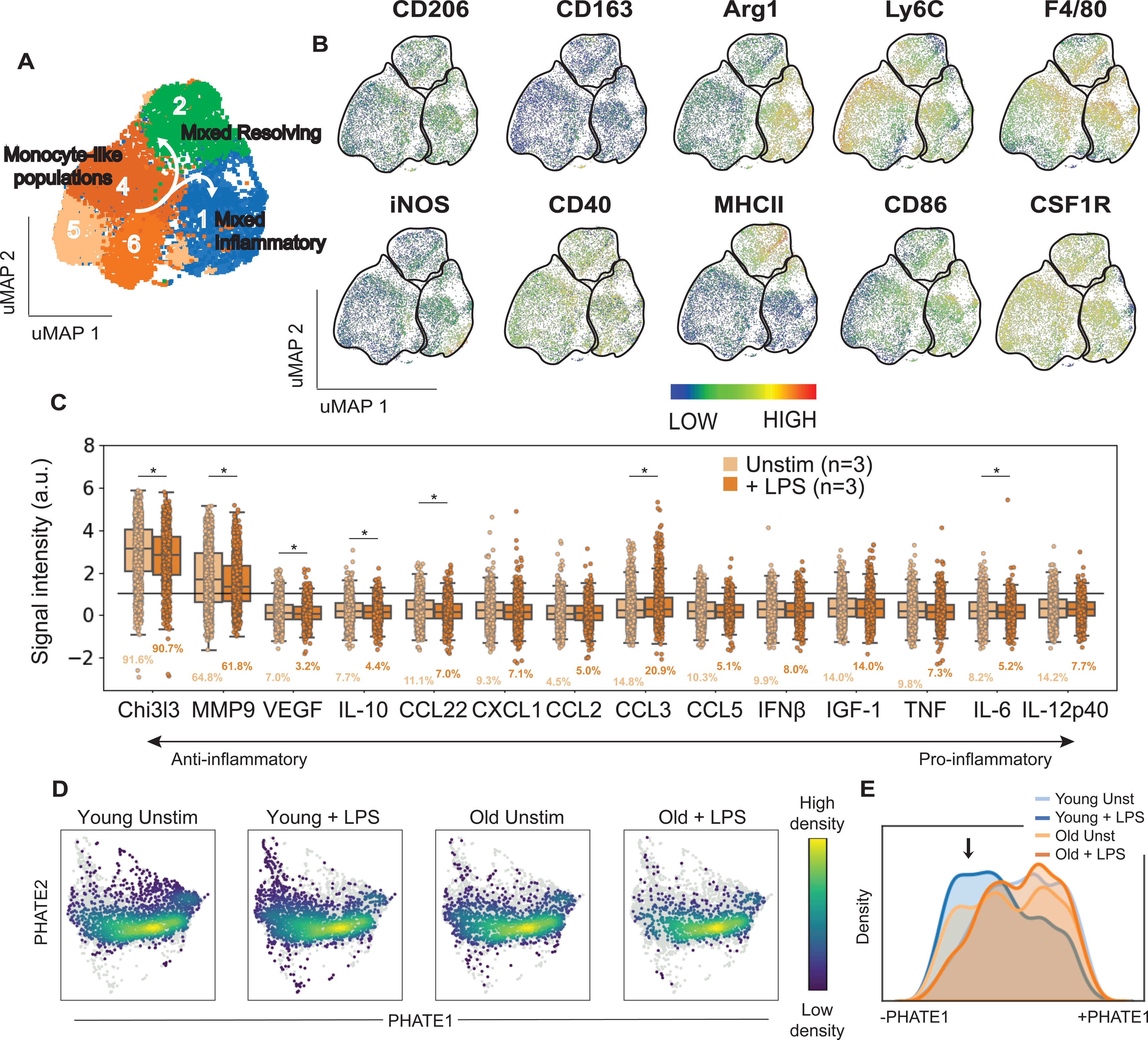
Aged macrophages display altered populations and inflammatory activation. A-B. UMAP and feature plots of flow cytometry data of 48-month-old macrophages isolated from naive skin and 24h wound beds. C. Box and whisker plots of single-cell secretion from F4/80+ macrophages from 24h wounds of 48-month-old mice treated with or without LPS. D. 2D PHATE visualization of dimensional single-cell secretion from F4/80+ macrophages isolated from 24h mouse wounds of aged mice either unstimulated or treated with LPS. Data include macrophages that secrete at least one measured cytokine above the threshold, colored by the kernel density estimation (KDE) for unstimulated or LPS-treated macrophages with grey indicating the other conditions. E. KDE for individual F4/80+ macrophages isolated from 24h wounds from aged mice with or without LPS stimulation along PHATE1 calculated from data in (D).

Next, we interrogated how aging impacts macrophage phenotypes via single cell secretomics of wound bed-derived macrophages from aged mice with and without ex vivo LPS stimulation. FACS purified CD45+, F4/80+, and CD11b+ macrophages from aged mice secreted high levels of resolving proteins, Chi3l3 and MMP9 (**Fig. 5C**), similar to wound bed macrophages from young mice (**Fig. 4C**). Aged macrophages also secreted IGF-1 but the fraction was somewhat lower than macrophages from young mice. Interestingly, LPS treatment of aged wound macrophages induced changes in the secretion of both resolving and inflammatory cytokines. Yet, unlike the wound bed macrophages from young mice (**Fig. 4C**), LPS significantly decreased secretion of Chi3l3, MMP9, VEGF, CCL22, and IL-6 in aged wound macrophages (**Fig. 5C**). Notably, unlike LPS-stimulated young wound macrophages (**Fig. 4C**), release of CXCL1, IGF-1, and IL-12p40 were not significantly altered with LPS stimulation in wound bed-derived macrophages from aged mice (**Fig. 5C**).

Combined PHATE visualization of the single-cell secretomics of the young and old macrophages revealed that, at baseline, aged macrophages somewhat resembled the secretome displayed by young macrophages (**Fig. 5D and 5E**). However, in response to LPS, aged macrophages did not significantly alter their secretion patterns (**Fig. 5D and 5E**). These data indicate that compared to young macrophages, age disrupts the secretory phenotypes of macrophages at baseline and in response to inflammatory stimuli.

### Aging rewires the metabolome of early wound bed macrophages

Age has been shown to impact metabolism so we hypothesized that the metabolism of aged wound macrophages would be changed compared to young wound macrophages (Bartke et al., 2003; Okuyama et al., 2021). To evaluate metabolic differences between early wound macrophages, we purified F4/80+ macrophages from young and old wounds 24h after injury using magnetic bead isolation and performed targeted LC/MS/MS metabolomics. We found 321 differentially abundant metabolites between the young and old groups (**Fig. 6A**). Principal component analysis (PCA) revealed distinct metabolite production of macrophages based on age (**Fig. 6B**). Pathway analysis revealed that old macrophages from early wounds harbored more metabolites in the TCA cycle, pentose phosphate pathway, and amino acid metabolism (**Fig. 6C and 6E**). In contrast, old macrophages from early wounds had fewer metabolites in beta-alanine, vitamin B6, and fatty acid metabolic pathways (palmitate and linoleic metabolism) (**Fig. 6D and 6F**), which broadly could represent a dysfunctional mitochondria phenotype since many of those pathways are restricted or related to this compartment (Bathina & Das, 2023).

**Figure 6.**
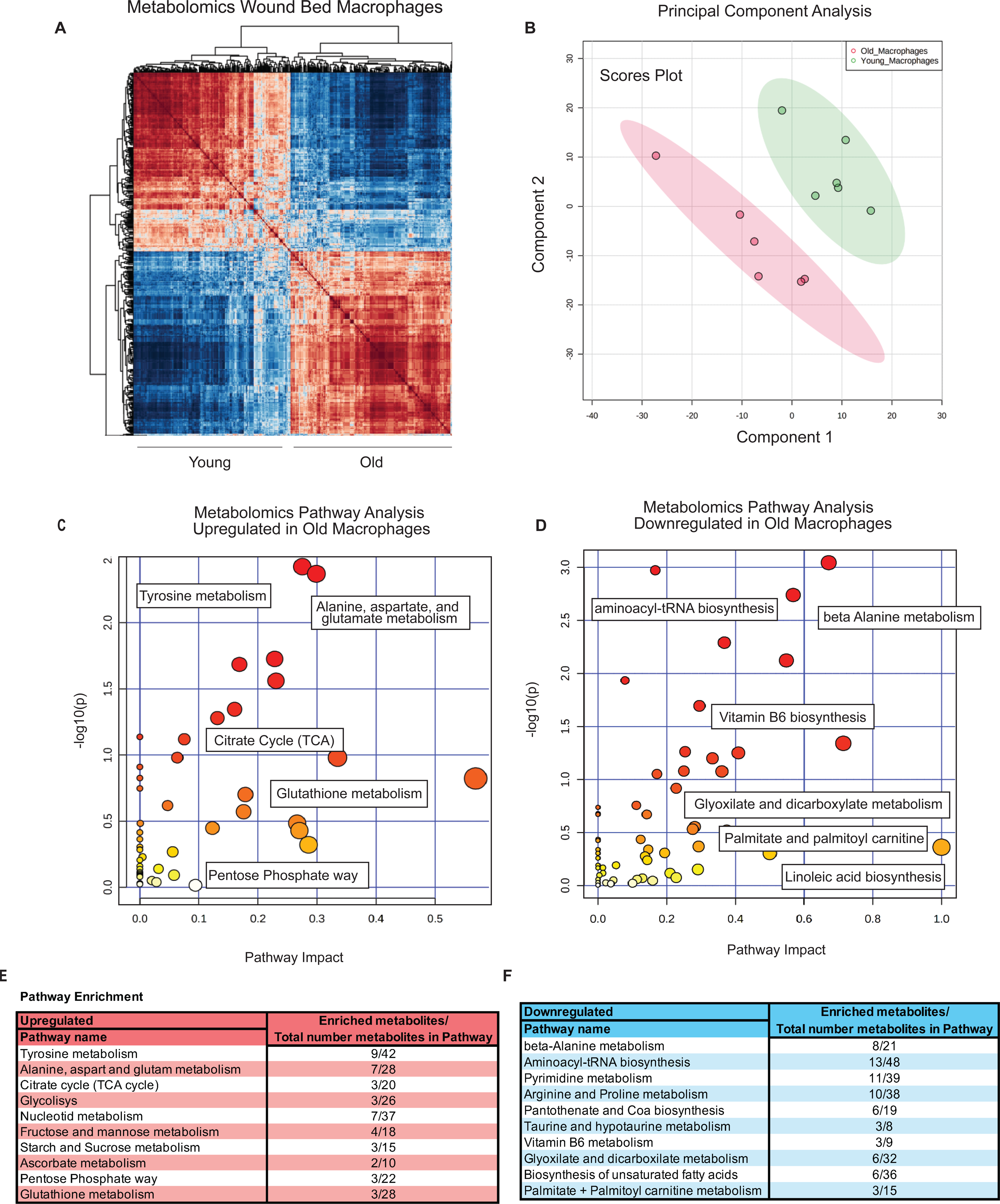
Aged macrophages alter their metabolic pathways. A-B. Heat map (A) and PCA plot (B) of metabolomic analysis of young vs old macrophages from 24h mosue wound beds. Data are pooled macrophages from 5 wound beds from each age. C-D. Pathway analysis plots of metabolic pathways upregulated (C) or downregulated (D) in 24h wound bed macrophages from aged mice. E-F. Tables of metabolic pathway analysis indicating enrichment scores of upregulated (C) or downregulated (D) pathways in metabolites from 24h wound bed macrophages from aged mice.

### IGF-1 and Chi3l3 can partially rescue the aging-associated changes in metabolism in macrophages

Since the secretome and metabolites within mitochondrial metabolism (TCA cycle, lipid metabolism) were altered in aged macrophages from early wounds (**Fig. 4-5**), we hypothesized that aged macrophages would display defects in the uptake of glucose and lipids and display altered mitochondrial mass compared to young macrophages in early wounds. We analyzed glucose uptake using the fluorescently labeled deoxyglucose analog, 2-(N-(7-Nitrobenz-2-oxa-1,3-diazol-4-yl)Amino (2-NBDG), and lipid uptake using fluorescent BODIPY-labeled fatty acids (Bala et al., 2021; Liao et al., 2005) (**Fig. 7A**). Analysis of Mitotracker green, which contains a mildly thiol-reactive chloromethyl moiety and accumulates in the mitochondrial matrix regardless of ΔΨm potential (Clutton et al., 2019), indicated mitochondrial mass (**Fig. 7A**).

**Figure 7.**
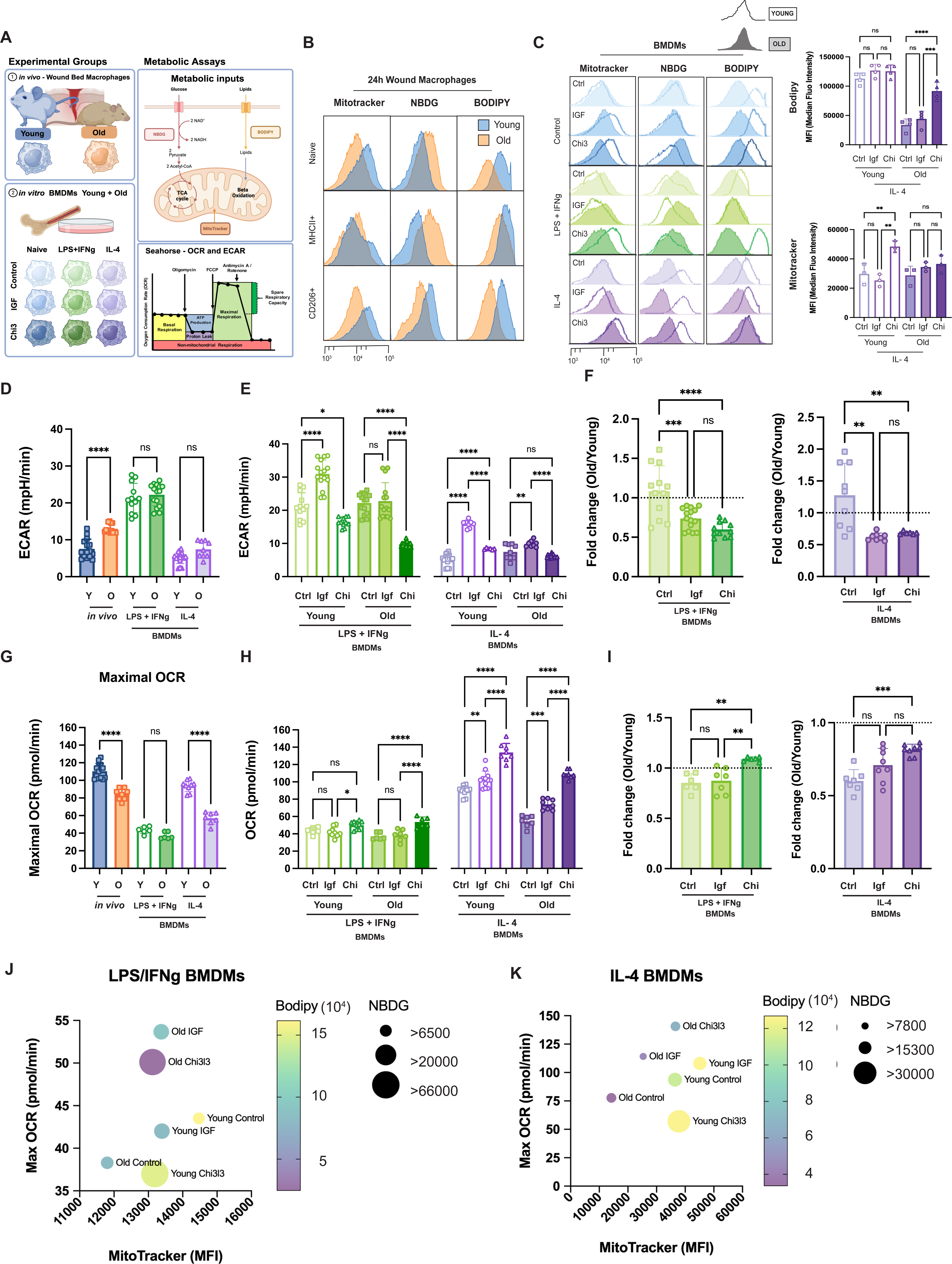
IGF and Chi3 are capable of partially rescuing the metabolic phenotypes of young and old BMDMs. A. Schematic illustrating the experimental groups and assays employed to investigate the role of age and IGF-1 (IGF) and Chi3l3 (Chi3) on macrophage metabolism. B. Histogram plots of FACS analysis of young (Y) and old (O) macrophages isolated from 24h wound beds and stained with deoxyglucose analog, 2-NBDG (NBDG), fluorescent lipid (Bodipy), and Mitotracker green. Data are representative of experiments from 3-6 mice. C. Histogram plots of FACS analysis of BMDMs from young (open) and old (filled) mice polarized as indicated, treated with vehicle (Ctrl), IGF-1 (Igf), or Chi3l3 (Chi), and stained with deoxyglucose analog, 2-NBDG (NBDG), fluorescent lipid (Bodipy), and Mitotracker green. Data are representative of experiments from 3-6 mice. D. Extracellular acidification rates (ECAR) of young (Y) and old (O) wound bed-derived macrophages (in vivo) and BMDMs polarized as indicated. Data are x experiments E-F. ECAR and fold change of young and old BMDMs polarized as indicated and treated with vehicle (Ctrl), IGF-1 (Igf), and Chi3l3 (Chi). G. Maximal Oxygen consumption rates (OCR) of young (Y) and old (O) wound bed-derived macrophages (in vivo) and BMDMs polarized as indicated. Data are x experiments H. Maximal OCR and fold change of young and old BMDMs polarized as indicated and treated with vehicle (Ctrl), IGF-1 (Igf), and Chi3l3 (Chi). J-K. Multiple variable plots of average NBDG, Mitotracker green, Bodipy, and Max OCR (shown in C and G) of young and old BMDMS polarized as indicated and treated with vehicle (Ctrl), IGF-1 (Igf), and Chi3l3. Data are combination of 4 to 6 independent experiments. Data in D-I are 4 to 6 mice (in vivo) and 4 independent experiments (BMDMs). Error bars indicate mean +/- SEM. p values calculated using one way ANOVA. ns, not significant.

Young wound-derived macrophages displayed less fluorescence for 2-NBDG, BODIPY, and Mitotracker than wound macrophages from old mice (**Fig. 7B**). Given the ability of polarization signals to alter metabolic inputs in BMDMs (Willenborg et al., 2021), we also analyzed how metabolic inputs and mitochondrial mass differed between wound-derived macrophages expressing more inflammatory or resolving proteins. Thus, we examined subsets of MHCII+ or CD206+ macrophages.

24h young wound-derived MHCII+ macrophages displayed less 2-NBDG, more BODIPY fluorescence, and a bimodal distribution of mitochondrial mass with most cells displaying lower mitochondrial mass than the entire population of wound-derived cells (**Fig. 7B, S6, and S8**). By contrast, 2-NBDG and BODIPY fluorescence was elevated in old MHCII+ macrophages with little change in mitochondrial mass compared to young macrophages expressing CD206 (**Fig. 7B**). Early wound CD206+ macrophages displayed increased 2-NBDG fluorescence in young mice and a slight decrease in aged mice compared to the bulk population (**Fig. 7B**). BODIPY and Mitotracker fluorescence were similar in CD206+ macrophages and decreased with age in this population (**Fig. 7B**). BMDMs isolated from aged mice and treated with and without polarization signals also displayed some age related changes in macromolecular uptake and mitochondria mass. These data indicate that aged wound macrophages display decreases in lipid uptake, increases in glucose uptake except in CD206+ wound macrophages, and largely reductions in mitochondrial mass except for a subset of MHCII+ wound macrophages.

Compared to young BMDMs with the same polarization induction, 2-NBDG fluorescence was significantly elevated in aged nonpolarized BMDMs, not changed in LPS/IFNg treated BMDMs, and significantly increased in IL-4 polarized BMDMs (**Fig. 7C and S7**). Mitotracker fluorescence was only significantly reduced in IL-4 polarized BMDMs with age compared to young BMDMs polarized with IL-4. (**Fig. 7C and S7**). Compared to young BMDMs with the same polarization induction, Bodipy fluorescence was significantly reduced in aged nonpolarized and IL-4-treated BMDMs, but not changed in LPS/IFNg treated BMDMs (**Fig. 7C and S7**). Thus, with some exceptions, aged BMDMs largely do not reflect changes in glucose and lipid uptake, or mitochondrial mass seen in aged skin wound macrophages.

Macrophage phenotype in young mice correlated with metabolic gene expression and increased secretion of IGF-1, which is a master regulator of global metabolism, aging, and mammalian longevity (Bartke et al., 2003; Okuyama et al., 2021). To determine if wound macrophage derived cytokines could alter these parameters in BMDMs, we treated BMDMs derived from young and old mice with IGF-1 (a more pro-inflammatory) and Chi3l3 (resolving), which were abundantly secreted by wound bed macrophages in young mice but altered with age (**Fig. 7A**). IGF-1 increased 2-NBDG fluorescence in nonpolarized, and IL-4 treated young BMDMs but not aged BMDMs and did not impact 2-NBDG fluorescence of either age in IFNg/LPS-treated BMDMs (Fig.7C and S8). IGF-1 did not significantly change Mitotracker fluorescence and only elevated Bodipy fluorescence in IL-4 treated BMDMs (**Fig. S7**).

By contrast, Mitotracker fluorescence was elevated by Chi3l3 treatment of young BMDMs in all polarization states (**Fig. 7C, 7D, and S7**) and was restored to young levels by Chi3l3 in aged IL-4 treated BMDMs (**Fig. 7D**). Chi3l3 also significantly increased Bodipy fluorescence in aged IL-4 treated BMDMs, and decreased Bodipy fluorescence in young non-polarized and IFNg/LPS-treated BMDMs (**Fig. S7**). Overall, these results suggest that Chi3l3 can increase mitochondrial mass of young polarized BMDMs and old IL-4 treated BMDMs, and can restore age-related reductions in lipid uptake in IL-4 treated cells (**Fig. 7C**).

To further investigate the metabolic characteristics of macrophages impacted by age, we measured extracellular acidification rate (ECAR, a measure of lactate production that reflects glycolysis rate) and oxygen consumption rate (OCR, which reflects the mitochondrial respiration of cells) with and without sequential inhibition of OXPHOS with oligomycin (Oligo) and rotenone/antimycin A (Rot/AA) to determine ATP and basal metabolic rates of oxygen consumption, respectively, in wound derived macrophages and BMDMs (**Fig. 7A**).

Consistent with elevated glucose uptake in wound macrophages with age (**Fig. 7C**), aged early wound-derived macrophages displayed elevated ECAR compared to young wound macrophages (**Fig. 7D),** but these age-related changes were not mimicked in polarized BMDMs from young and old mice (**Fig. 7D**). Basal OCR is decreased in old wound-bed derived macrophages and in inflammatory LPS/IFNg polarized BMDMs and consistently increased in IL-4 treated BMDMs, independent of age. (**Fig. S6A**).

ATP-production OCR was not different between young and old 24-hour wound-derived macrophages but was significantly reduced in aged BMDMs polarized with either IFNg/LPS or IL-4 (**Fig. 7G**). Maximal OCR was reduced considerably in aged 24h wound-derived macrophages and IL-4 treated BMDMs but not IFNg/LPS treated BMDMs (**Fig. 7J**). These data reveal several points: 1) age shifts the metabolism of wound-derived macrophages toward glycolysis and reduces mitochondrial metabolism and 2) BMDMs stimulated with IL-4 display glycolytic and maximal OXPHOS similar to wound-derived macrophages.

Next, we analyzed whether IGF-1 or Chi3l3 could rewire ECAR and OCR in BMDMs (**Fig. 7A-7C)**. We treated polarized BMDMs from young mice with IGF-1 and Chi3l3 and performed metabolic analysis with Seahorse. Both IGF-1 and Chi3l3 significantly increased ECAR in both IL-4-treated BMDMs, but only IGF-1 increased ECAR in LPS/IFNg-stimulated BMDMs (**Fig. 7F**). IGF-1 reduced basal OCR in LPS/IFNg, and IL-4 stimulated BMDMs, and increased maximal OCR in IL-4-treated BMDMs. By contrast, Chi3l3 elevated basal and ATP production OCR in LPS/IFNg treated BMDMs and all three OCR measurements in IL-4 treated BMDMs (**Figs. 7I, 7L, and S6C**). Thus, Chi3l3 can rewire the metabolism of BMDMs to mimic the high mitochondrial metabolism found in wound-derived macrophages of young mice.

Next, we analyzed whether IGF-1 or Chi3l3 could restore the metabolic defects in aged BMDMs. While IGF-1 did not significantly impact ECAR or OCR measurements in young vs old polarized BMDMs, we found that Chi3l3 significantly reduced ECAR and increased the ATP-production OCR and maximal OCR of both LPS/IFNg treated and IL-4-treated BMDMs (**Fig. 7F, 7I, and 7L**). Importantly, the impact of Chi3l3 on maximal OCR levels was almost to that of young BMDM levels, showcasing the ability of this cytokine to increase mitochondrial activity (**Fig. 7E**). Visualization of the average measurements for macromolecule uptake (2-NBDG and BODIPY fluorescence), mitochondrial mass (Mitotracker fluorescence), and maximal OCR highlights the ability of Chi3l3 and IGF-1 to elevate mitochondrial mass and metabolism in old BMDMs of both inflammatory and resolving phenotypes to young levels (**Figs. 7J and 7K**).

## Discussion

Our study highlights the complexity of macrophage populations in early wound environments, offering insights into their transcriptional and metabolic diversity. Rather than adhering to the traditional binary models of macrophage activation often described in vitro, we find that macrophages within early skin wounds co-express mRNAs and secrete proteins for both inflammatory and resolving functions. Heterogeneity has been observed for macrophages, including those resident within tissues (Chakarov et al., 2019; Heieis et al., 2023), within the tumor microenvironment (Loyher et al., 2018), and between early and late-stage wounds in the skin (Willenborg et al., 2021). While the presence of macrophages expressing resolving proteins such as CD301b has been observed in early wounds (Shook et al., 2016; Wasko et al., 2022), our data indicate that individual early wound macrophages display both inflammatory and resolving phenotypes with a continuum of gene expression. We observe that monocytes form two mixed populations of macrophages within early skin wounds, which is consistent with trajectory analyses of macrophage gene expression in colorectal (Ji et al., 2024) and breast cancer (Azizi et al., 2018). Such hybrid phenotypes reflect the growing understanding that macrophages function as integrators of diverse environmental cues, transitioning between states that are not easily classified (Guilliams & Scott, 2022; Papachristoforou & Ramachandran, 2022).

Our data highlight the ability of macrophage-derived cytokines to shape their phenotypes in mouse skin wounds. An emerging theme is the ability of local microenvironments to produce specialized macrophage functions in many tissues, including the skin (Lavin et al., 2014; Okabe & Medzhitov, 2014). However, the signals that drive these microenvironments have largely remained elusive. Microbial-derived nucleic acids and proteins, and intracellular DNA induce inflammatory macrophage phenotypes in vivo through activation of pattern recognition receptors like Toll-like receptors, retinoic acid-inducible gene I (RIG-I)-like receptors, or scavenger receptors (Rasheed & Rayner, 2021). Other immune cells, such as T cells and eosinophils, can instruct macrophage phenotype via cytokine release (Rasheed & Rayner, 2021). In the liver, local lipid exposure activates macrophages (Guilliams & Scott, 2022). In contrast, retinoic acid and other local signals can activate Gata6 in peritoneal macrophages to induce macrophage compartmentalization and B cell IgA production (Jayakumar et al., 2022; Shi et al., 2024). Local tissue signals may also regulate resolution as lipid mediators and efferocytosis of specific cell types may influence macrophage tissue repair phenotypes (Spadaro et al., 2017).

Here, we functionally link Chi3l3, which is associated with resolving macrophages (Ponomarev et al., 2007), to the regulation of macrophage metabolism in macrophages in early mouse skin wounds. Chi3l3 is an enzymatically inactive chitinase that can promote antiviral immunity (Osborne et al., 2014a) and neutrophil accumulation via γδ T cells during helminth infection (Osborne et al., 2014a, 2014b) and Th2 polarization of T cells in vitro (Kim et al., 2018). During neuroinflammation, Chi3l3 can also promote oligodendrocyte differentiation via epidermal growth factor receptor activation (Starossom et al., 2019). Whether Chi3l3 acts via EGFR or other signaling pathways to modulate macrophage metabolism will be an exciting area of future research.

Emerging evidence supports the importance of the tissue niche in controlling macrophage metabolism (Heieis et al., 2023). In vitro studies have highlighted the ability of macrophages to rewire their metabolism via bioenergetics, nutrient uptake, and metabolic pathways, depending on their inflammatory or resolving phenotype (O’Neill & Pearce, 2016; van den Hoogen et al., 2020; Wculek et al., 2022). A recent, elegant study in mice revealed that early-stage wound macrophages do not use glycolytic metabolism but rely on oxidative phosphorylation for ROS production and inflammatory function (Willenborg et al., 2021), contrary to in vitro polarization models (O’Neill & Pearce, 2016). Our data are consistent with the use of mitochondrial metabolism of early wound macrophages and previous *in vitro* and *in vivo* studies showing that IL-4 stimulates OXPHOS and glucose uptake and metabolism in macrophages (Huang et al., 2016; Willenborg et al., 2021). We further demonstrate that aged macrophages of early wounds are more glycolytic and that IGF-1 and Chi3l3 can drive metabolic rewiring of in vitro polarized BMDMs and aged wound macrophages.

Our data in skin wound healing is consistent with other studies showing that inflammaging, or chronic inflammation associated with aging, is characterized by macrophages with defective activation profiles. While the mechanisms that drive inflammaging are not well understood (van Beek et al., 2019; Vu et al., 2022), metabolic rewiring of aged macrophages has been implicated in several studies. Prior studies have shown that aged human monocyte-derived macrophages (Minhas et al., 2019) and in specific clusters of mouse skin wound macrophages (Vu et al., 2022) have a metabolic shift from OXPHOS to glycolysis. Furthermore, changes in macrophage signaling in aged wounds supports age-inducing inflammatory phenotypes via IL-1 signaling (Vu et al., 2022). Rewiring of metabolism can reverse macrophage aging phenotypes via prostaglandin E2 signaling in the brain, supporting an essential role for metabolism in driving aging phenotypes (Minhas et al., 2021). In adipose tissue, inflammaging via elevated adipocyte secretion of the matricellular protein, secreted protein acidic and rich in cysteine (SPARC) also acts in part through activation of glycolysis (Ryu et al., 2022). Our data bolster the growing theme of macrophage driven defects with age to highlight their autocrine/paracrine role in the wound bed niche to drive age-dependent metabolic modulation during Inflammaging. Given the impaired ability of tissue repair to occur in aged skin and other tissues (Wilkinson & Hardman, 2020), these data and future investigations have implications for therapeutic avenues for pathological conditions like age-related wounds and other disorders, where disrupted macrophage responses can profoundly affect tissue homeostasis and repair.

## Material and Methods

### EXPERIMENTAL MODEL AND SUBJECT DETAILS

#### Mice

Wild-type C57BL/6J mice (males, young 2-4 months and old 48 months) were purchased from Jackson Laboratories. All mice were housed in the Yale Animal Resources Center in specific pathogen-free conditions. All animal experiments were performed according to the approved protocols of the Yale University Institutional Animal Care and Use Committee.

#### Cultured Cells

Bone marrow-derived macrophages (BMDMs) were generated as previously described (Trouplin et al., 2013). In brief, a syringe extracted bone marrow from the tibias and femurs of mice’s hind legs flushing media. Afterward, red blood cells were lysed with ammonium-chloride-potassium lysis buffer (Lonza). The cells were incubated at 37C with 5% CO2 in a culture-treated Petri-dish with BMDM media (RPMI supplemented with 10% FBS, 100 U/mL penicillin, 100 mg/mL streptomycin, 1% sodium pyruvate, 25 mM HEPES buffer, 2mM L-glutamine, and 50 mM 2-mercaptoethanol). After 4 hours, the nonadherent cells were transferred to a new non-tissue culture-treated Petri dish and incubated with BMDM media supplemented with 20 ng/mL M-CSF (PeproTech). After seven days, BMDMs were polarized cells stimulated with LPS (Invivogen) and IFNg or IL-4 (Peprotech).

#### Fabrication of antibody barcodes and microchamber array chips for single-cell secretion

Antibody barcodes and microchamber array chips were fabricated as previously described (13). Briefly, chips were manufactured with PDMS (RTV615, Momentive, parts A and B in a 10:1 ratio) from silicon masters through soft lithography techniques. To fabricate antibody barcodes, a poly-L-lysine microarray glass slide (Erie Scientific) was bound to the PDMS chip designed for flow patterning, and 2 ml of each antibody was then flowed through individual microchannels until dry. Antibody pairs used in this study are listed in Table S3. The SCBC used in this study contains 3080 rectangular chambers with the dimensions 35 × 35 × 1850 mm (width × depth × length).

#### Microwell assay for single-cell secretion profiling

Single-cell secretion profiling experiments were performed as previously described (Lu et al., 2013; Xue et al., 2015). Briefly, capture antibodies (Key resources table) were flow patterned onto epoxy silane-coated glass slides (Super-Chip; ThermoFisher). The polydimethylsiloxane (PDMS) microwell arrays and antibody barcode glass slides were blocked using complete RPMI. BMDMs were suspended in complete RPMI supplemented with ten ng/mL M-CSF, added to the PDMS microwell array, and allowed to adhere overnight. The next day, BMDMs were stimulated as indicated with complete RPMI supplemented with 125 nM of live cell marker (Calcein AM; ThermoFisher) to allow automatic live cell detection. The BMDMs in the PDMS microwell array were then covered with the antibody barcode slide, secured with plates and screws, and allowed to incubate for 8 hours. At the end of the incubation period, the device was imaged with an automated inverted microscope (Axio Observer; ZEISS) to record the sound position and cell locations. The device was then disassembled, and the sandwich immunoassay was performed: The glass slide was incubated with a mixture of detection antibodies (Key resources table) for 2 hours, followed by incubation with 20 mg/mL streptavidin-APC (eBioscience) for 30 minutes, rinsed with PBS and deionized water, and scanned with a Genepix 4200A scanner (Molecular Devices).

#### Flow Cytometry

BMDMs were lifted in ice-cold PBS+EDTA with gentle scraping and immediately fixed with Fix buffer (BD Biosciences) for 10 minutes at 37C. Cells were then transferred to a 96-well u-bottom plate. After washing, cells were blocked with Fc receptor antibody (eBioscience, CD16/32, 1:200 dilution) on ice for 15 mins in FACS buffer (PBS + 2% FBS). Cells were stained with antibodies in 200 uL for 1 hour at room temperature. All samples were acquired on an Attune NxT Flow Cytometer (Gabby, please confirm) and analyzed with FlowJo (FlowJo, LLC).

#### Fluorescence-activated cell sorting

Mouse back skin and wound beds were dissected and digested into single cells using 1:100 collagenase 1A (Worthington) and Liberase TM for macrophage subsets, as previously described (26, 41, 72). Cells were resuspended in FACS staining buffer (1% BSA in PBS with two mM EDTA). To examine immune cell subsets, cells were stained with the following antibodies for 30 minutes on ice: CD45-PECy7 (eBioscience, 1:2000), CD11b-Alexa700 (eBioscience, 1:500), F4/80-eFluor450 (eBioscience, 1:50), CD206-Alexa488 (Biolegend, 1:500), MHC-II-eFluor450 (eBioscience, 1:1000), CD64-PerCp (Biolegend, 1:500) and CD301b-Alexa660 (eBioscience, 1:100). Wound macrophages were defined as CD45+; CD11b+; F4/80+ cells. Sytox Orange or Sytox Blue (Invitrogen, 1:1000) was added immediately before sorting using a FACS Aria III with FACS DiVA software (BD Biosciences) to exclude dead cells. Cells were sorted into 10% fetal bovine serum (FBS) in RPMI, and flow cytometry analysis was performed using FlowJo Software (FlowJo).

#### RNA extraction and Real-Time PCR

Wound bed immune cells and BMDMs were lyzed using TRIzol LS (Invitrogen). RNA was extracted from the aqueous phase using the RNeasy Plus Micro Kit (Qiagen). cDNA was generated using one ug of total RNA with the Superscript III First-Strand Synthesis Kit (Invitrogen) per manufacturer instructions. All quantitative real-time PCR was performed using SYBR green on a Light Cycler 480 (Roche). Primers for specific genes are listed in Supplementary Table 2. Results were normalized to β-actin HMBS and HPRT as previously described (Ling & Salvaterra, 2011).

#### Single-cell RNA Sequencing

Single-cell RNA sequencing was performed as described (Justynski et al., 2023). Briefly, scRNA-seq data from 24-and 48 hr mouse wound beds were processed using the standard cellranger pipeline (10X Genomics). Downstream analysis was performed using the Scanpy package in Python (Buttner et al., 2019). Cells were filtered for quality control to avoid doublets and dead cells. Dimensionality reduction and downstream data visualization were completed using the Scanpy implementation of UMAP (Lim & Qiu, 2023) and the ShinyCell package in R (Ouyang et al., 2021), respectively. Data is presented as scaled log-normalized mRNA counts (i.e., expression). Differentially expressed genes (DEGs) across time points were calculated using the rank_genes_groups function from the Scanpy module in Python with default parameters.

#### Seahorse quantification of Extracellular acidification rate (ECAR) and Oxygen Consumption Rate (OCR)

OCRs were measured in primary cultures using an XF96 Bioanalyzer (Seahorse Bioscience) as described previously (Forni et al., 2016; Forni et al., 2017). An assay medium composed of 114 mM NaCl, 4.7 mM KCl, 1.2 mM KH^2^PO_4_, 1.16 mM MgSO_4_, 2.5 mM CaCl_2_ (pH 7.2), and 10 mM glucose was used. Cells were seeded in an XF 96-well microplate at 100,000 cells/well in 100 μL of growth medium and incubated overnight at 37°C in a humidified atmosphere of 95% air and 5% CO_2_. Before the assay, the medium was replaced with 100 μL assay medium. Cells were preincubated for 1 hr at 37°C without CO_2_. The first 3 measurements without any treatment were taken as the Basal OCR. ATP production-linked O_2_ consumption was determined by the addition of oligomycin (0.5 μg/mL). After 3 measurement cycles, 5 μM of the uncoupler CCCP was added to determine the maximal respiratory capacity. After a further 3 cycles of measurement, 1 μM rotenone and 1 μM antimycin A were added, ablating mitochondrial O_2_ consumption. All respiratory modulators were used at concentrations determined through preliminary titration and experiments to determine the ideal cell density were performed.

#### Metabolomics data preparation and analysis

Metabolomics was performed at the Yale Chemical Metabolism Core (CMC). In brief, cells were quenched by 150 µL of ice-cold quenching buffer. Quenching buffer recipe was as follows: 20% MEOH into 0.1% formic acid, 3 mmol/L NaF, and 5.5 μg/mL D8-Phenylalanine (used as an internal standard). Then, the material was transferred to a LC/MS/MS V-bottom plate on dry ice, stored in –80°C freezer until liquid is completely frozen and lyophilized overnight. The lyophilized powder was reconstituted in 50 µL/well D4-taurine solution (another internal standard). Six biological replicates were created in each scenario. Each sample was run with two columns and two MS modes per column to get the best coverage of metabolites: reverse phase column with positive and negative MS mode; hypercarb column with positive and negative MS mode. The targeted group of metabolites was based on our in-house IROA libraries (∼600 metabolites in total) and the untargeted curation was based on the Kyoto Encyclopedia of Genes and Genomes (KEGG) database (∼2,700 metabolites). To call a metabolite significant, the same rules were applied as in the gene expression data analysis.

### QUANTIFICATION AND STATISTICAL ANALYSIS

Unless otherwise specified, data were analyzed using GraphPad Prism for Mac (GraphPad Software, La Jolla, CA) with significance set at p < 0.05. To determine significance between two groups, comparisons were made using Student’s t-test, while multiple groups were compared using a one-way ANOVA with Bonferroni’s post hoc using GraphPad Prism for Mac.

#### Single-cell secretion profiling and data processing

Device images were analyzed using a custom script in MATLAB (MathWorks) to detect healthy locations automatically and the number of cells per well, extract all signals from each well and process the data (https://github.com/Miller-JensenLab/Single-Cell-Analysis). In brief, after automatic well and live cell detection, signal image registration, and manual curation, the software automatically extracted the intensity signal from each antibody for all the microwells in the device. Each antibody’s signal across the chip was normalized by subtracting a moving Gaussian curve fitted to the local zero-cell well intensity values. A secretion threshold for each antibody was set at the 99th percentile of all normalized zero-cell wells. Data was transformed using the inverse hyperbolic sine with cofactor set at 0.8x secretion threshold. Statistics Data were presented as means ± SEM unless otherwise specified. Statistical analysis was performed by ordinary 2-way ANOVA and the Dunnett method to correct multiple comparisons as defined in the figure legends. All analyses were performed using Prism 8.4.1 software (GraphPad). For single-cell distributions, statistics were performed using a bootstrapping procedure to calculate the confidence intervals associated with sampling error in single-cell data. To obtain confidence intervals through bootstrapping, the single-cell datasets for each condition were sampled 10,000 times with replacement, and the metric of interest was calculated for each resampled dataset. We then calculated a 95% confidence interval for these resampled datasets, and statistical significance was assigned to pairwise comparisons with non-overlapping confidence intervals. This bootstrapping procedure was done using custom scripts in MATLAB and Python.

#### Multidimensional PHATE analysis

The dimensionality reduction algorithm known as the potential of heat diffusion for affinity-based transition embedding (PHATE) was used further to visualize high-dimensional secretion data from the microwell assay. PHATE analysis was performed using MATLAB and Python packages from the Krishnaswamy lab (https://github.com/KrishnaswamyLab/PHATE). Standard parameters were used to make 2-dimensional PHATE projections from the nine and 10-dimensional BMDM datasets. Manual adjustments were made to the t parameter for optimal visualization. PHATE plots were colored by kernel density estimation (KDE) using Gaussian kernels or by relative secretion levels of each protein as indicated; cells below the secretion threshold were greyed out for visualization. Extracted PHATE parameters were analyzed using custom software written in Python. KDE plots of the PHATE 1 coordinates were plotted using Seaborn. We fitted a Gaussian mixture model with three components to the distribution of PHATE 1 coordinates. We chose three components based on minimizing the Akaike information criteria (AIC) and Bayesian information criteria (BIC) scores.

## Acknowledgments

The authors would like to thank all the Horsley and Miller-Jensen laboratory members for their feedback and critical analysis of the manuscript. We would also like to thank the Yale Animal Resources Center (YARC) staff for animal husbandry, the Yale School of Medicine and Yale Science Hill Flow Cytometry Core (especially), and the Yale Center for Genome Analysis, which receives funding from NIH Shared Equipment grant #1S10OD028669-01. M.F.F. received funding from a Pew Fellowship. V.H. is funded by NIH NIAMS R01s AR076938, AR0695505, and AR075412. KM-J is funded by NIH U01-CA238728, R01-CA238728, and R01-GM123011.

## Author contribution

Conceptualization: MFF, KMJ, VH

Methodology: MFF, KMJ, VH

Software: GP, KB, MFF

Validation, Analysis and Investigation: MFF, GP, KB, WK, AA, YX, OJ, AG, NCS, KMJ, VH

Resources: KMJ, VH

Data Curation: MFF, GP, KB, KMJ, VH

Writing – Original Draft MFF and VH

Writing – Review & Editing all authors

Visualization MFF, GP, KB, KMJ, VH

Supervision KMJ, VH

Funding Acquisition KMJ, VH

---

**Supplementary Figure 1.**
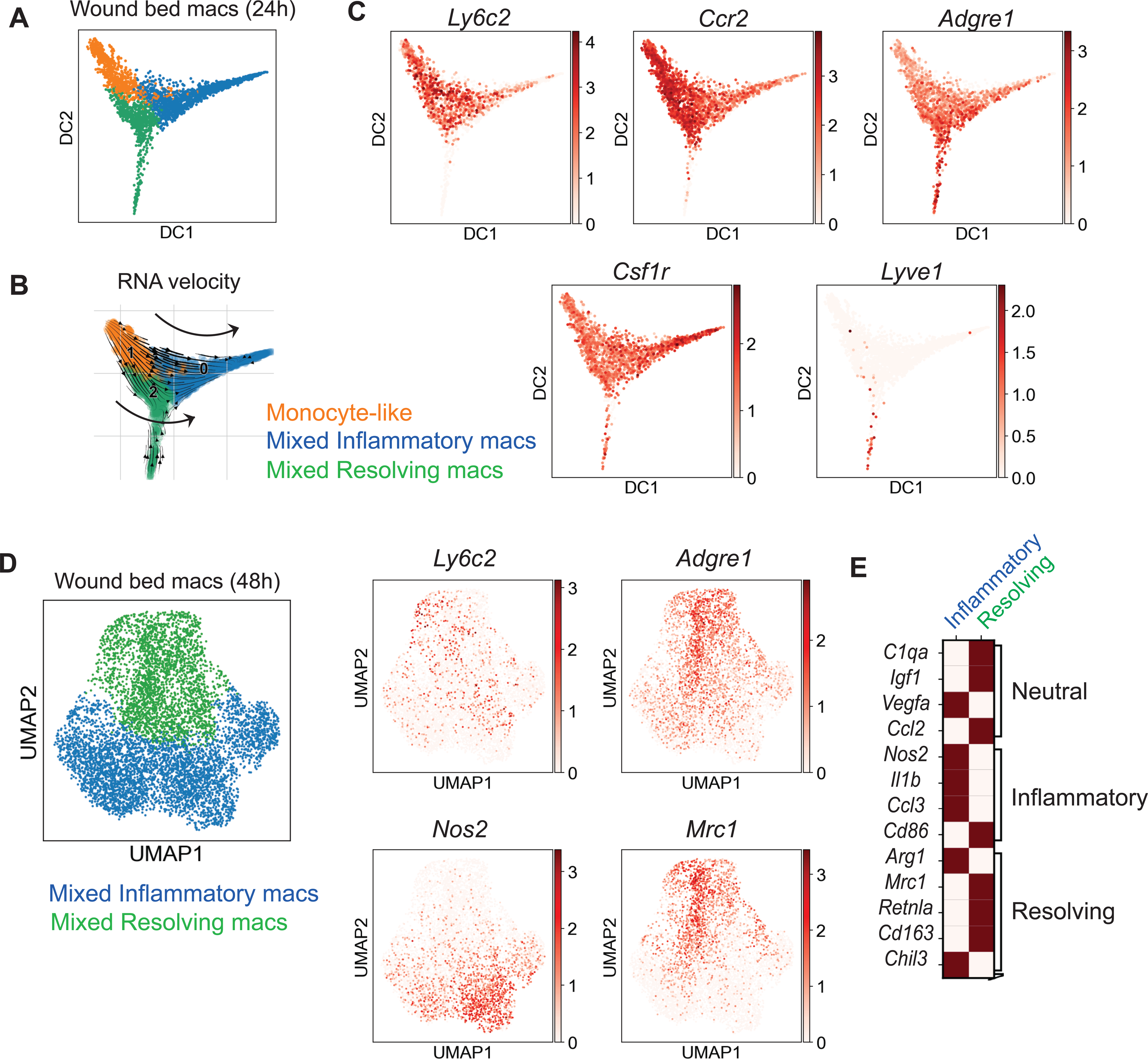
Related to Figure 1. A. Differential Composition plots of scRNA sequencing data of wound bed macrophages and monocytes. Data are from GSE223660. B. Trajectory analysis of scRNA sequencing data of wound bed macrophages and monocytes. Data are from GSE223660. C. Feature plots of scRNA sequencing data of wound bed macrophages and monocytes. D. UMAP and feature plots of scRNA sequencing data from 48-hour (h) macrophages (macs). E. Heatmap of mRNAs associated with mixed inflammatory and resolving macrophages in 48h wound bed macrophage scRNA sequencing data. F. Feature plots of single cell RNA sequencing data of bone marrow derived macrophages.

**Supplementary Figure 2.**
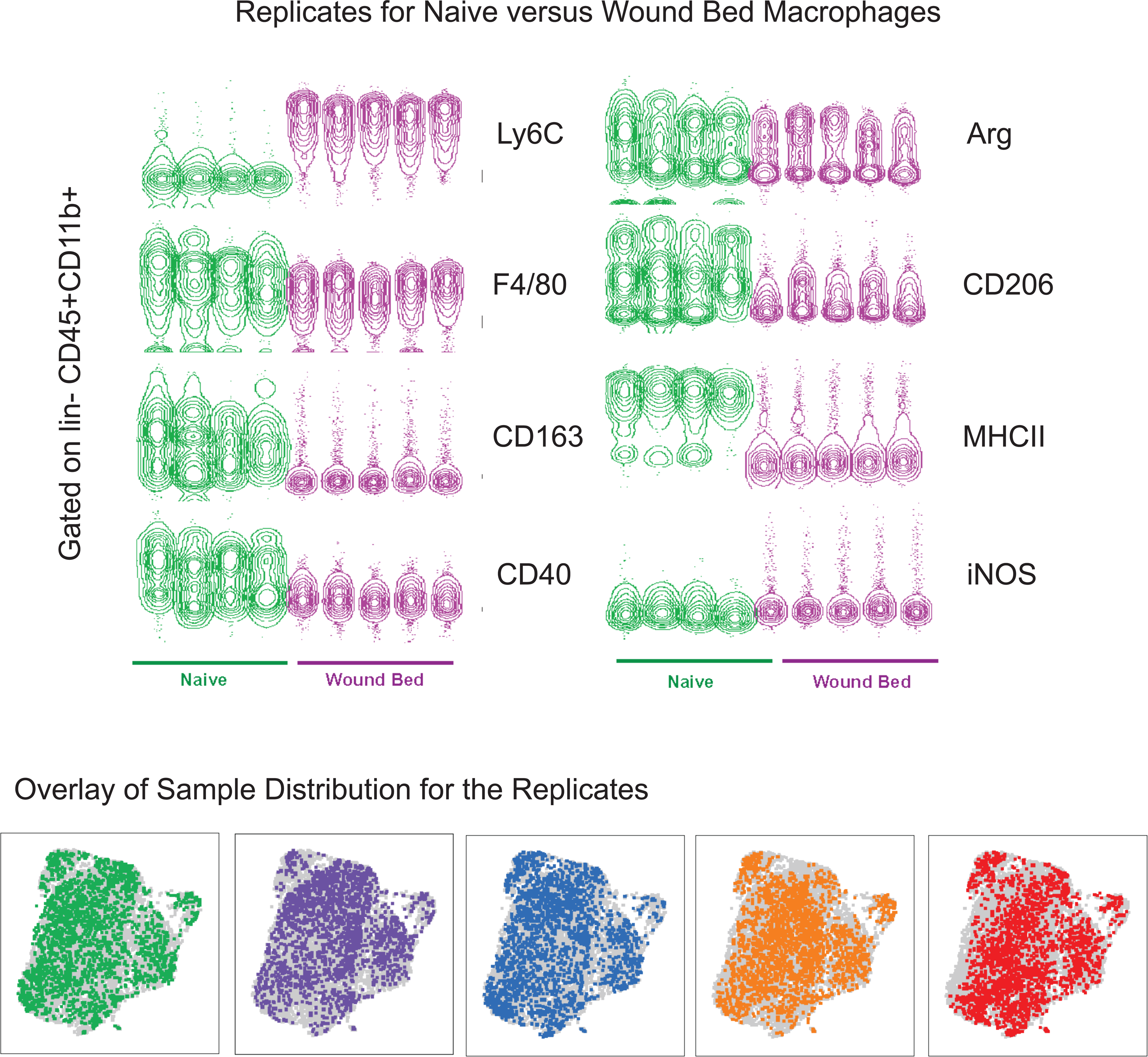
Related to Figure 3. A. Individual replicates for naïve vs. wound bed flow cytometry. 4 mice from each group. B. Individual distribution from markers for wound bed macrophages flow cytometry.

**Supplementary Figure 3.**
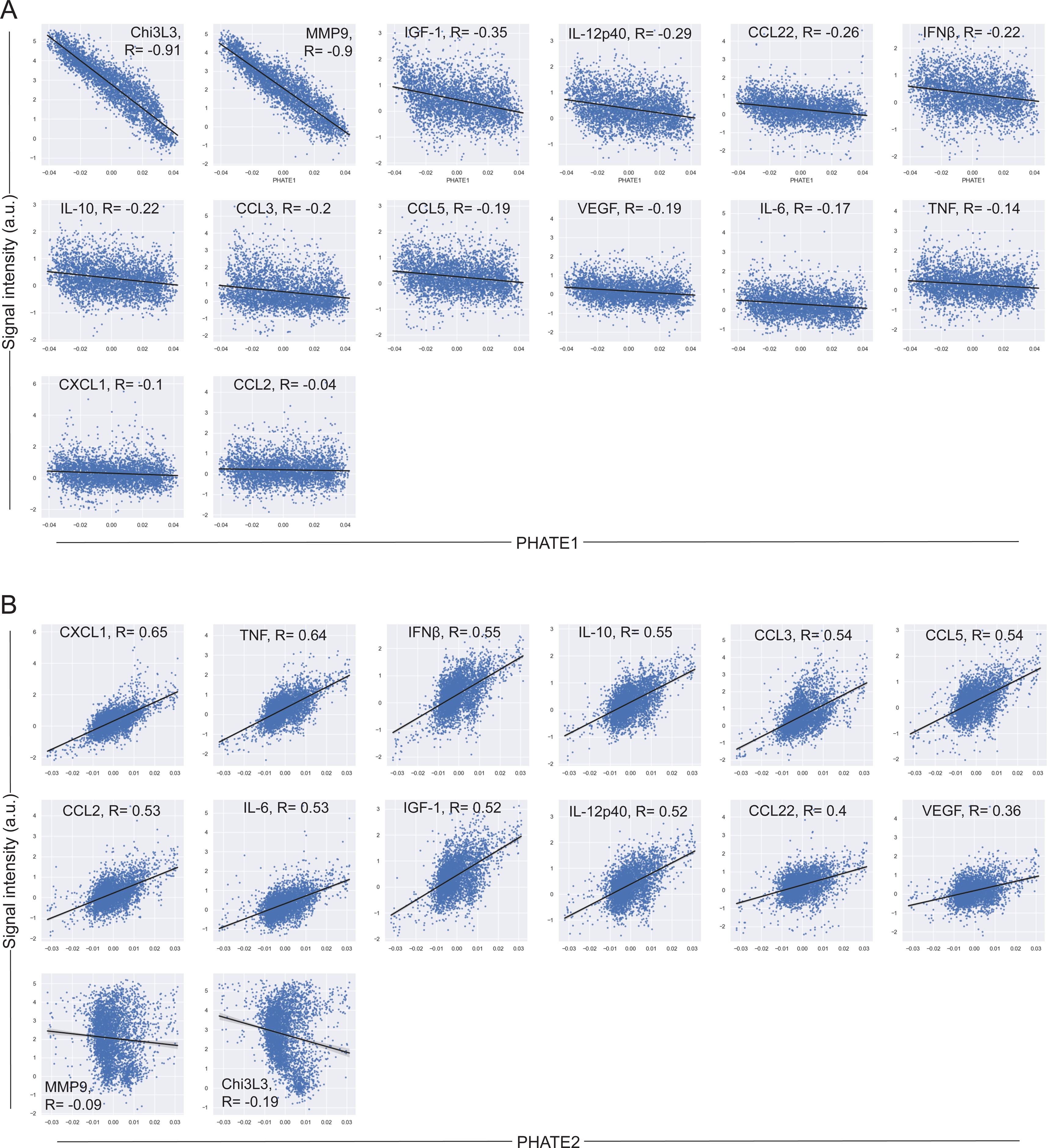
Related to Figure 4. Replicates for individual wound beds for high dimensional flow cytometry for macrophages isolated from 24h skin wounds.

**Supplementary Figure 4.**
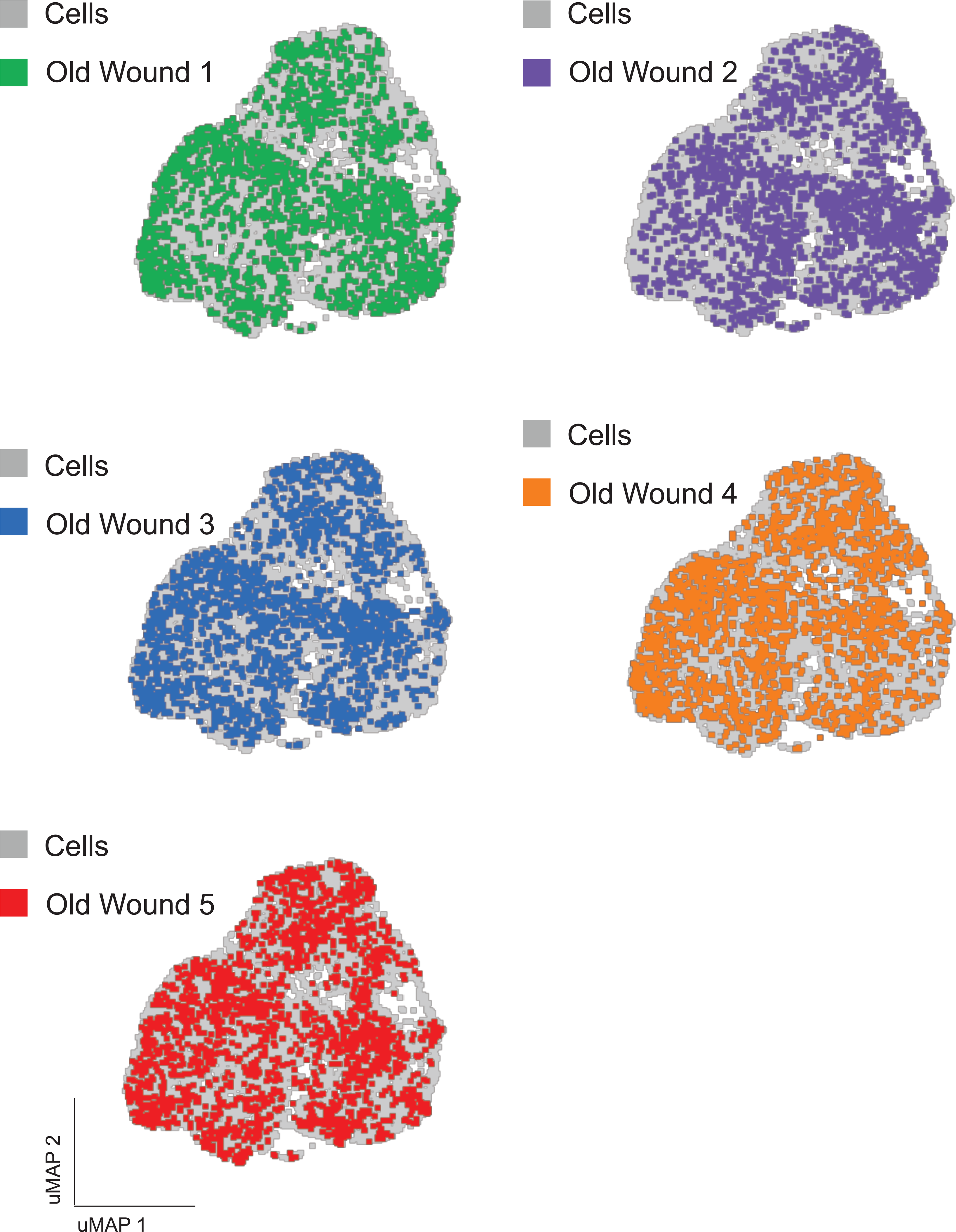
Related to Figure 5. Cytokine secretion intensity in the microwell device correlates with PHATE axes (PHATE1 (A) and PHATE 2 (B). Scatter plots show correlation between single-cell secretion for the indicated cytokines from wound bed macrophages. Coordinates extracted from two-dimensional PHATE embeddings of the data. Linear best fit lines are shown in black, and the plots are ordered by Pearson correlation coefficient (shown as R).

**Supplementary Figure 5.**
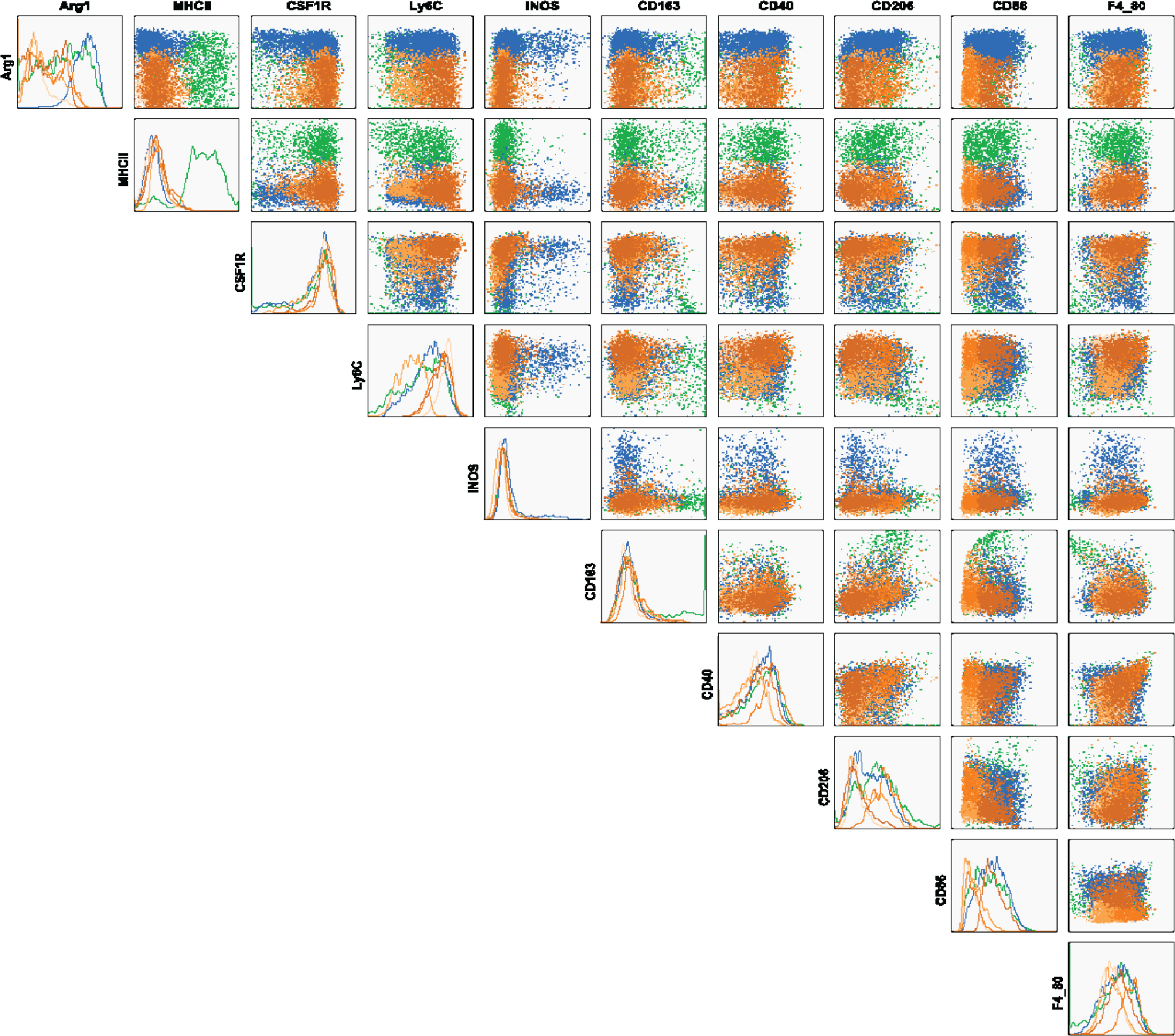
Related to Figure 2 and 4. Individual populations of FACS markers (Arg1, MHCII, CSF1R, Ly6C, iNOS, CD163, CD40, CD206, CD86, F4/80) for high dimensional flow cytometry of young wound bed. Identified clusters are labeled in each plot.

**Supplementary Figure 6.**
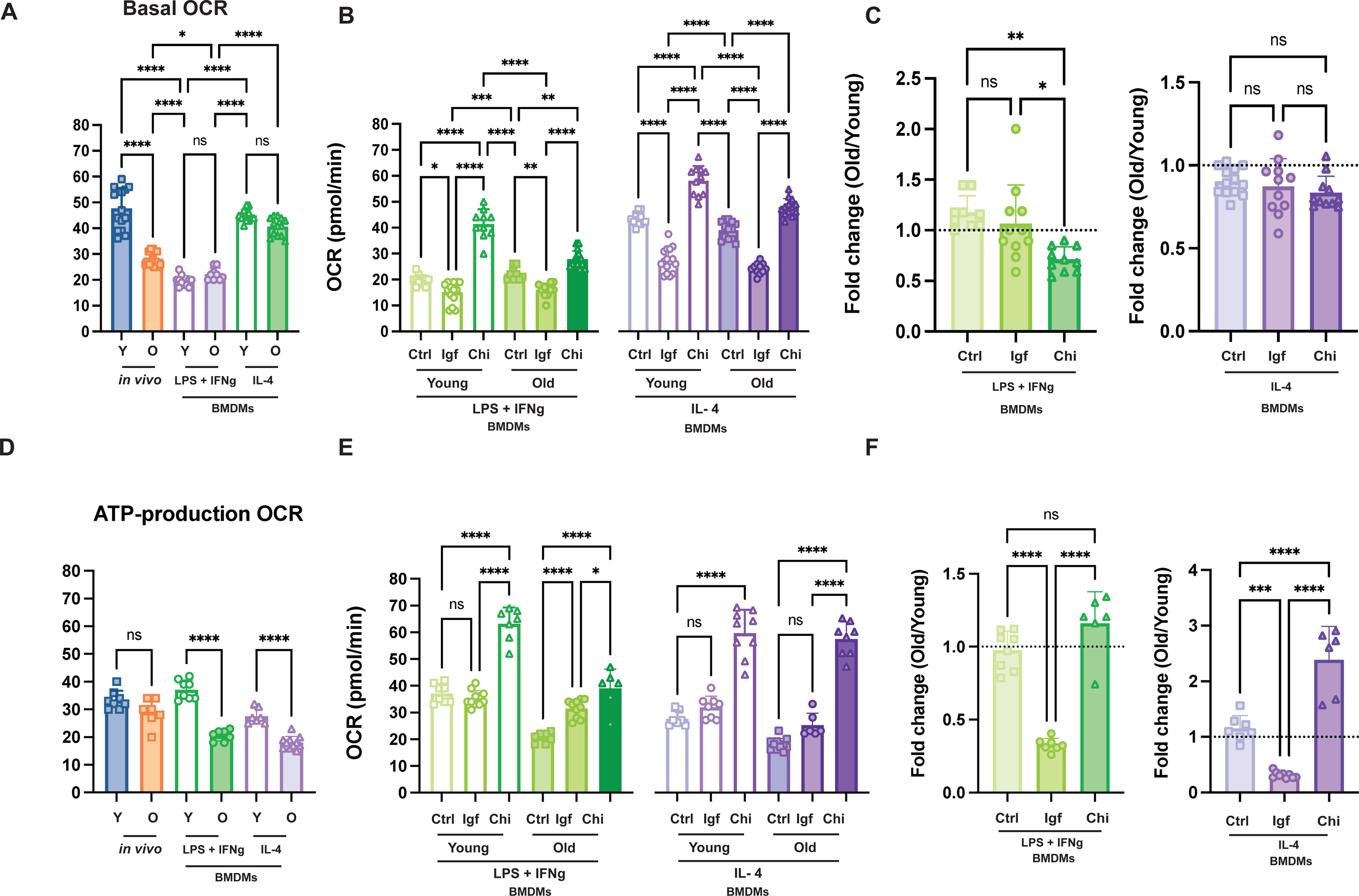
Related to Figure 7. A. Basal oxygen consumption rates (OCR) of young (Y) and old (O) wound bed-derived macrophages in vivo and polarized BMDMs as indicated. B-C. Basal OCR and fold change of young and old BMDMs polarized as indicated and treated with vehicle (Ctrl), IGF-1 (Igf), and Chi3l3 (Chi). D. ATP-production related OCR of young (Y) and old (O) wound bed-derived macrophages in vivo and BMDMs polarized as indicated. E-F. ATP-production related OCR and fold change of young and old BMDMs polarized as indicated and treated with vehicle (Ctrl), IGF-1 (Igf), and Chi3l3 (Chi).

**Supplementary Figure 7.**
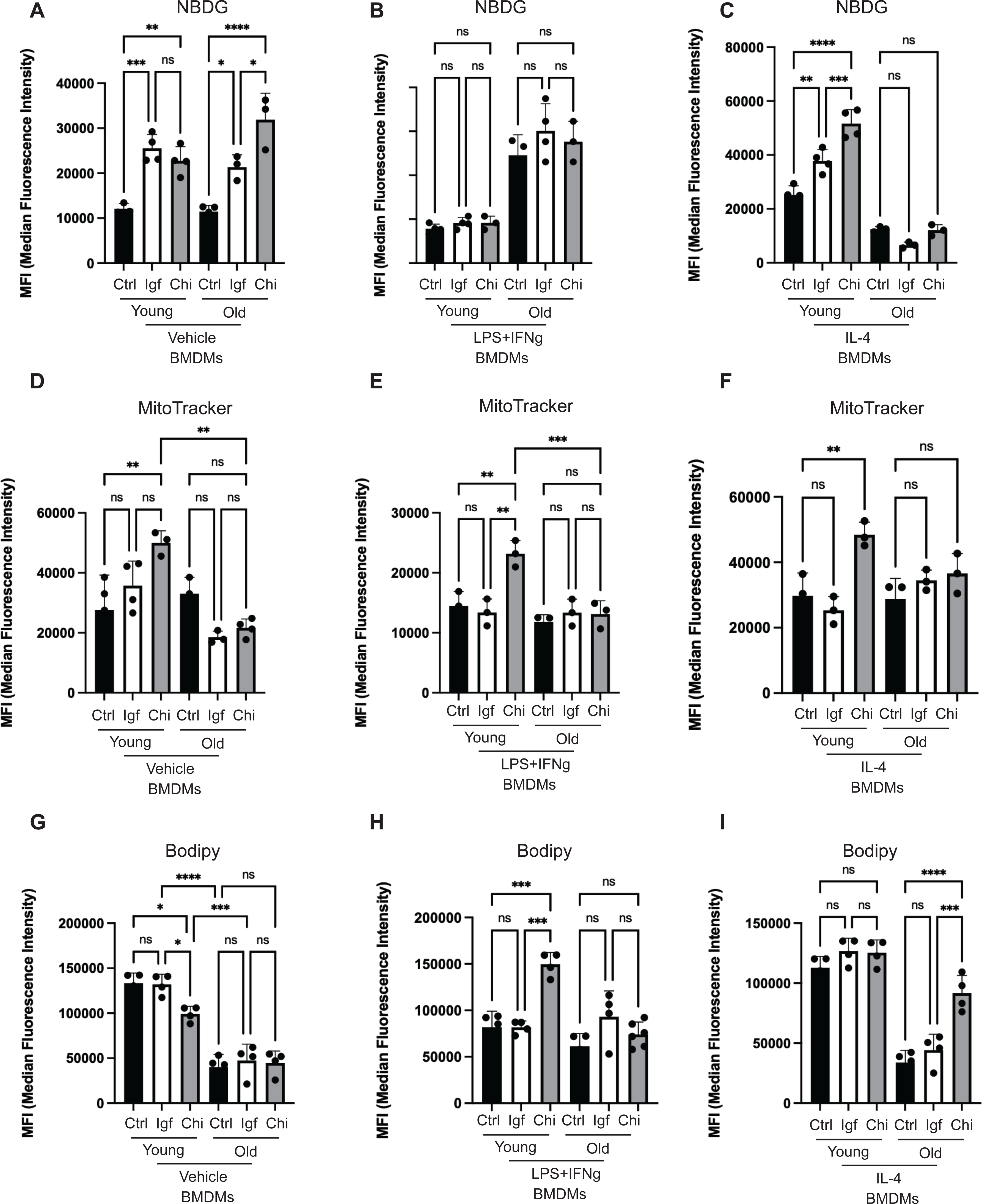
Related to Figure 7. Bar plots of <u>M</u>edium <u>F</u>luorescence Intensity (MFI) of FACS analysis of young (Y) and old (O) macrophages isolated from 24-hour wound beds and bone marrow derived macrophages (BMDMs) from young and old mice polarized as indicated (LPS+IFNg or IL-4), treated with vehicle (Ctrl), IGF, or Chi3l3. Data are representative of experiments from 3-6 mice. A. Unpolarized BMDMs and wound bed macrophages stained with deoxyglucose analog, 2-NBDG (NBDG). B. LPS+IFNg polarized BMDMs stained with deoxyglucose analog, 2-NBDG (NBDG). C. IL-4 polarized BMDMs stained with deoxyglucose analog, 2-NBDG (NBDG). D. Unpolarized BMDMs and wound bed macrophages stained with and Mitotracker green. E. LPS+IFNg polarized BMDMs stained with and Mitotracker green. F. IL-4 polarized BMDMs stained with and Mitotracker green. G. Unpolarized BMDMs and wound bed macrophages stained with fluorescent lipid (Bodipy). H. LPS+IFNg polarized BMDMs stained with fluorescent lipid (Bodipy). I. IL-4 polarized BMDMs stained with fluorescent lipid (Bodipy).

**Supplementary Figure 8.**
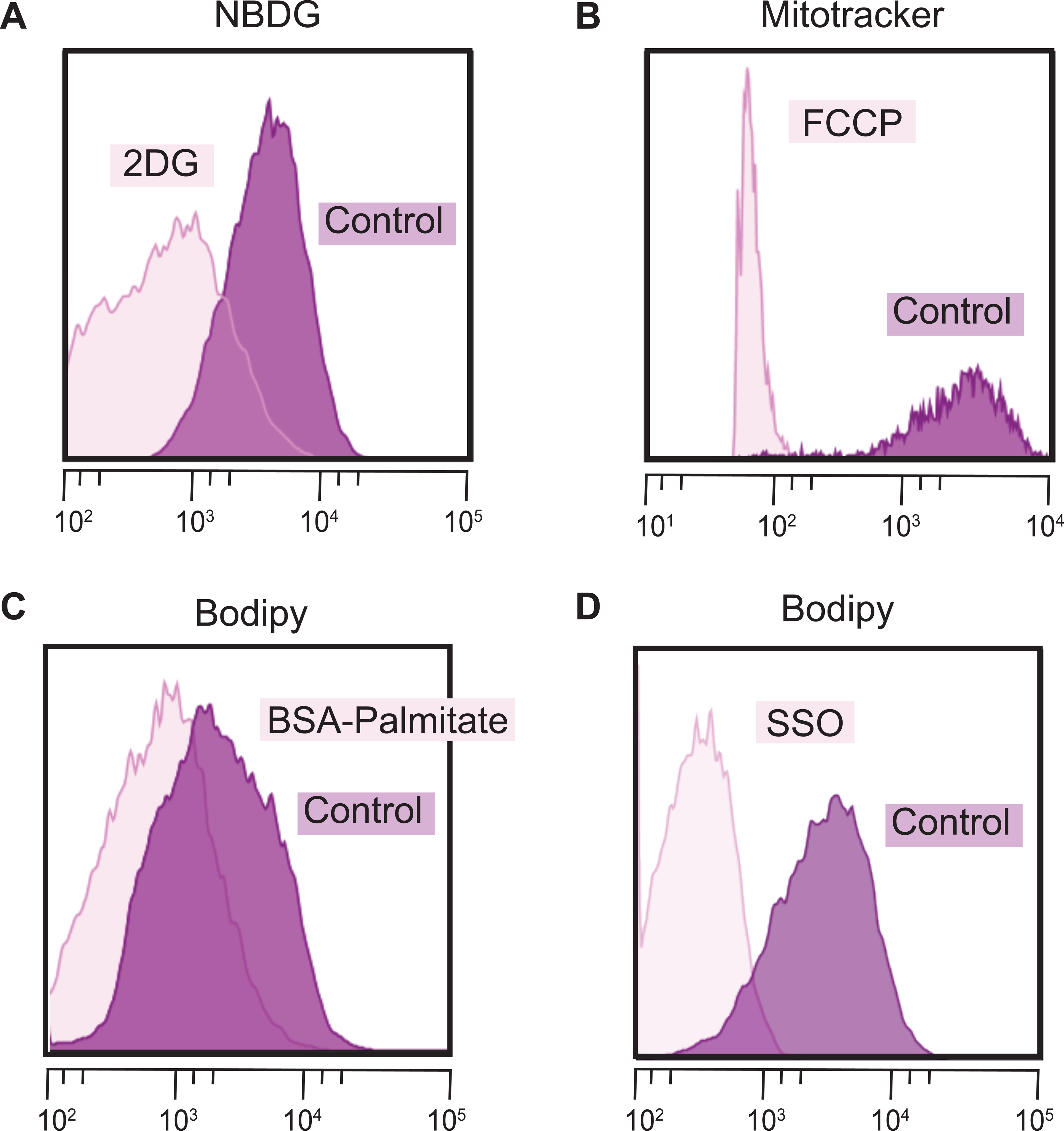
Related to Figure 7. Histogram plots of FACS analysis of young bone marrow derived macrophages (BMDMs) stained with deoxyglucose analog, 2-NBDG (NBDG) (A), Mitotracker green (B), or fluorescent lipid (Bodipy) (C-D) and treated with vehicle (Control) or appropriate negative control: 2-deoxyglucose, an inhibitor of glucose uptake (2DG), carbonyl cyanide-p-trifluoromethoxy phenylhydrazone a mitochondrial uncoupler (FCCP), BSA-Palmitate (6:1), Sulfo-N-succinimidyl oleate (SSO), which blocks the fatty acid receptor/transporter CD36.

